# The forensic landscape and the population genetic analyses of Hainan Li based on massively parallel sequencing DNA profiling

**DOI:** 10.1101/2020.03.27.011064

**Authors:** Haoliang Fan, Zhengming Du, Fenfen Wang, Xiao Wang, Shao-Qing Wen, Lingxiang Wang, Panxin Du, Hai Liu, Shengping Cao, Zhenming Luo, Bingbing Han, Peiyu Huang, Bofeng Zhu, Pingming Qiu

**Affiliations:** School of Forensic Medicine, Southern Medical University, Guangzhou 510515, China; School of Basic Medicine and Life Science, Hainan Medical University, Haikou 571199, China; First Clinical Medical College, Hainan Medical University, Haikou 571199, China; Department of Psychiatry, The First Clinical Medical College, Shanxi Medical University, Taiyuan 030001, China; Institute of Archaeological Science, Fudan University, Shanghai 200433, China; MOE Key Laboratory of Contemporary Anthropology and B&R International Joint Laboratory for Eurasian Anthropology, School of Life Sciences, Fudan University, 200438 Shanghai, China; Division of the Criminal Investigation Department, Henan Provincial Public Security Bureau, Zhengzhou 450003, China; School of Traditional Chinese Medicine, Hainan Medical University, Haikou 571199, China

**Author notes:** Corresponding authors. (Qiu PM) and (Zhu BF).

**Keywords:** Hainan Li, forensic landscape, population genetics, ForenSeq^™^, DNA Signature Prep Kit, Massively parallel sequencing

## Abstract

Due to the formation of the Qiongzhou Strait by climate change and marine transition, Hainan island isolated from the mainland southern China during the Last Glacial Maximum. Hainan island, located at the southernmost part of China and separated from the Leizhou Peninsula by the Qiongzhou Strait, laid on one of the modern human northward migration routes from Southeast Asia to East Asia. The Hlai-language speaking Li minority, the second largest population after Han Chinese in Hainan island, is the direct descendants of the initial migrants in Hainan island and has unique ethnic properties and derived characteristics, however, the forensic associated studies on Hainan Li population are still insufficient.

Hence, 136 Hainan Li individuals were genotyped in this study using the MPS-based ForenSeq^™^ DNA Signature Prep Kit (DNA Primer Set A) to characterize the forensic genetic polymorphism landscape, and DNA profiles were obtained from 152 different molecular genetic markers (27 autosomal STRs, 24 Y-STRs, 7 X-STRs, and 94 iiSNPs). A total of 419 distinct length variants and 586 repeat sequence sub-variants, with 31 novel alleles (at 17 loci), were identified across the 58 STR loci from the DNA profiles of Hainan Li population. We evaluated the forensic characteristics and efficiencies of DAPA, it demonstrated that the STRs and iiSNPs in DAPA were highly polymorphic in Hainan Li population and could be employed in forensic applications. In addition, we set up three Datasets, which included the genetic data of (I). iiSNPs (27 populations, 2640 individuals), (II). Y-STRs (42 populations, 8281 individuals), and (III). Y-haplogroups (123 populations, 4837 individuals) along with the population ancestries and language families, to perform population genetic analyses separately from different perspectives.

In conclusion, the phylogenetic analyses indicated that Hainan Li, with a southern East Asia origin and Tai-Kadai language-speaking language, is an isolated population relatively. But the genetic pool of Hainan Li influenced by the limited gene flows from other Tai-Kadai populations and Hainan populations. Furthermore, the establishment of isolated population models will be beneficial to clarify the exquisite population structures and develop specific genetic markers for subpopulations in forensic genetic fields.

## 1. Introduction

Hainan province, which is made up of 35.4 thousand square kilometers land areas (including Hainan island, Xisha, Zhongsha, and Nansha archipelagoes in the South China Sea) and about 2 million square kilometers sea lands, is located at the southernmost tip of the People’s Republic of China and separated from the Leizhou Peninsula to the north by the Qiongzhou Strait. While, Hainan island, as the dominating part of Hainan with 1,823 kilometers coastlines and 33,900 km^2^ areas (18°10’~20°10’N, 108°37’~111°03’E), is the second largest island after Taiwan island in China [1–5]. (The specific geographical location of Hainan province is shown in Fig. 1.) According to the Hainan Statistical Yearbook 2019 which was presented on the website of The People’s Government of Hainan Province (http://www.hainan.gov.cn/), the permanent resident population was 9.343 million people in the year of 2018, including 9.251 million people who were the household registered population. On account of the unique geographical location and population expansion historically [4–9], nowadays, Hainan island has been taken sharp in the specific and unparalleled treasure-house assembling ethnology, nation science, and linguistics with more than thirty ethnic minorities (consisting of Han, 81.89%; Li, 16.37%; Miao, 0.87%; Zhuang, 0.44%; Hui, 0.15%; and others, 0.28%) and a wide range of languages, including Chinese language (Hainanese, Danzhou Dialect, Tanka Dialect, Hakka Chinese, Mandarin), Tai-Kadai language (Hlai, Ong Be, and Cun language), Hmong-Mien language (Miao or Hmong language), Austronesian family/Malayo-Polynesian language (Tast language) [1–3, 10–19].

**Fig. 1.**
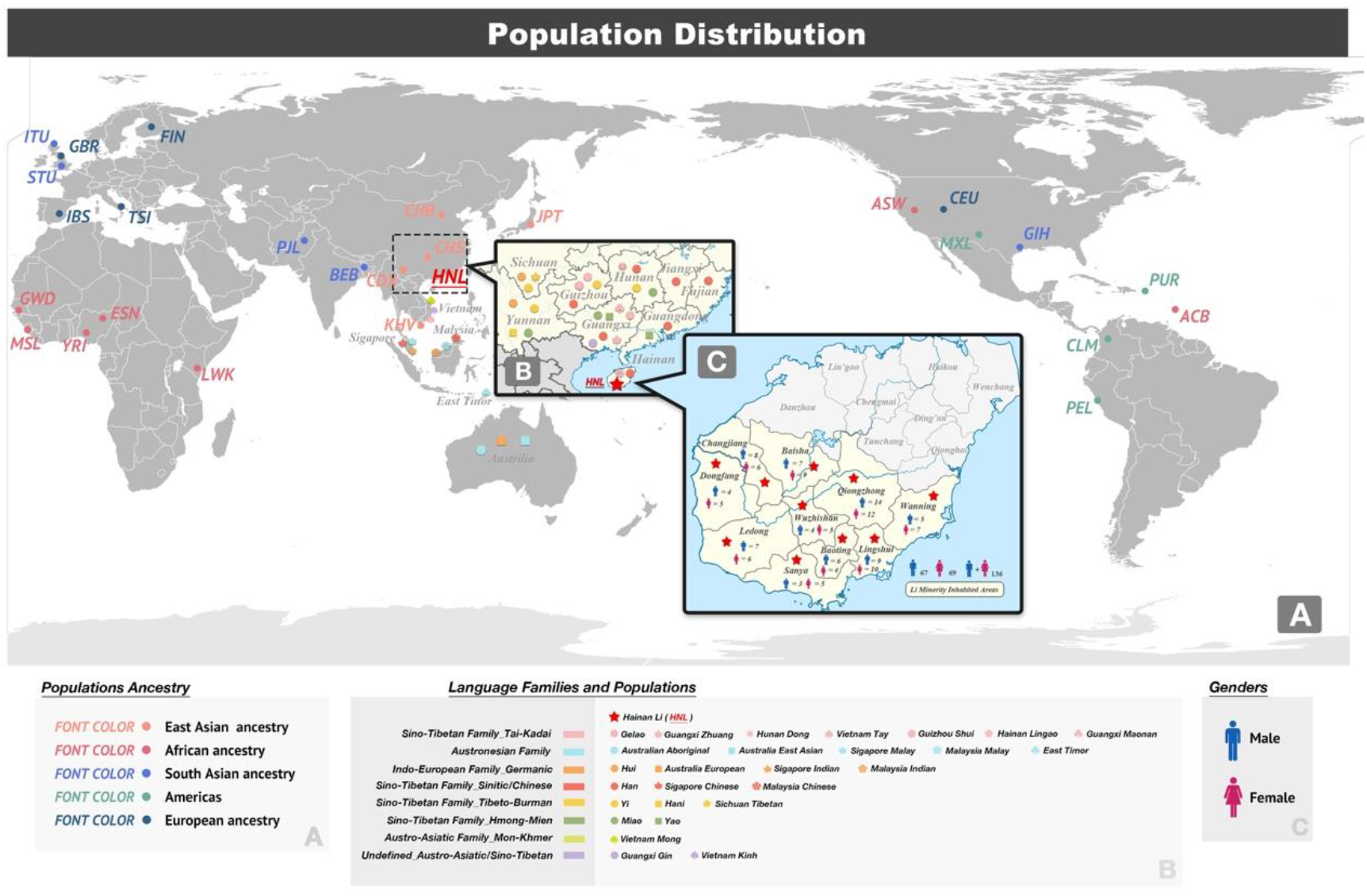
Geographic position and population distribution. **A.** 1 KG populations. **B.** Populations in Dataset II. **C.** Sampling positions of Hainan Li.

The initial peopling of Hainan island, where laid on one of the modern human northward migration routes from Southeast Asia to East Asia, was likely to the early Holocene and/or even the late Upper Pleistocene (around 7 - 27 thousand years ago, kya) based on complete mtDNA genomes sequencing [15], when Hainan island was connected with the mainland southern China and/or northern Vietnam, and the sea level was around 80-120 meter below present day [6, 15, 20, 21]. While, the archaeological findings in Luobi Cave of Sanya County (~10kya) and Xinjie Shell-heap site of Dongfang (~6kya) in Hainan island suggested that the definite traces of anatomically modern humans in Hainan island were evidenced and the colonization and human activities settled on Hainan island could be traced back to the Upper Paleolithic Period (~10kya) or the Upper Pleistocene (1-12.6kya) [22–26]. Due to the formation of the Qiongzhou Strait by the abrupt rise in sea level and the exact evidences which could be traced back to the Neolithic Period [4, 6–8, 21, 27, 28], Hainan island isolated from the mainland southern China during the Last Glacial Maximum (~14.5kya) and the colonization of Hainan island by the ancestors of the Li people, the direct descendants of the original migrants, could be at least traced back to the transition period between the Upper Paleolithic Period and the Early Neolithic Period (~ 10kya) [10–12, 15, 17–19, 24]. From the past to the present, as the primitive tribes, Li population still maintains the ancient cultures of the Neolithic Period, the old tattoo and divination cultures (Jí Bŭ) inherited from the ancient times to this day and age, and the old lifestyles, bark cloth and original ceramic making [1–4, 10], also preserved until current time. Li population mainly distributed in the whole region of Hainan island in ancient times, while, the mass landings of the Han (since BC 214 in Qin Dynasty) and Lingao people (about BC 500 during the Spring and Autumn period) compressed the living space of Hainan Li, making Li from the north to the south of Hainan island [1, 2, 9, 29, 30]. (Nowadays, the main distributions of Hainan Li minority are shown in Fig. 1C.) In all ages, ever since the primordial settlements in Hainan island, Hlai language was the dominant language for the original migrants and their descendants, and the intangible cultural and spiritual heritages within Hainan Li were preserved by oral transmission without ancient writings to hand down. Hlai, which is one important branch of the Tai-Kadai language, is widely spoken by more than 1.51 million Hainan Li for daily communications at present [31, 32].

In the field of massively parallel sequence (MPS), for the past decade, there are tremendous advances had been made in the aspect of accuracy, speed, read length, and throughput, as well as the substantial cost reduction. Together, these progresses democratized this technology and laid a solid foundation for the future development of abundant applications in basic subjects and translational research areas, such as clinical diagnostics, agrigenomics, and forensic sciences [33–39]. The ForenSeq^™^ DNA Signature Prep Kit, as the first commercial MPS-based and forensic related kit formally released in 2015, made full use of the MPS method’s advantages by means of the simultaneous and compound amplification of either greater than 150 loci including various types of genetic markers. The large-scale performances and functional tests had been widely validated for the ForenSeq^™^ DNA Signature Prep Kit, the DNA Primer Set A (DPMA, 152 loci totally: 27 common, forensic autosomal STRs, 24 Y-STRs, 7 X-STRs, and 94 identity informative SNPs) and the DNA Primer Set B (in total 230 loci: DMPA plus 22 phenotypic informative SNPs and 56 biogeographical ancestry SNPs), which also include robustness, reproducibility, species specificity, consistency and sensitivity of detection [40–47]. In addition, the applicability of the DNA Signature Prep Kit had been evaluated and verified on challenging samples, DNA mixture, degradative DNA [48–50], formalin-fixed paraffin-embedded tissues [49, 51], and remains of skeletons and bones [48, 52–54], to extend the applications for forensic sciences [55–68]. Hence, on the basis of the sufficient validations of the stability and homogeneity, more essential data of worldwide populations are needed urgently at present to improve the standard experimental procedures, promote the efficiencies of MPS-based system, and strengthen and unify the contacts between capillary electrophoresis-based (CE-based) and MPS-based data for the basic and extended applications of forensic sciences and others.

As the aborigines in Hainan island where was the entrances to East Asia (either the southern entrance from Indo-China peninsula or the northern entrance from Central Asia) [12, 69–71], Hainan Li, the second largest population after Han Chinese in Hainan island, has unique ethnic properties and derived characteristics which serves as potential “transfer station” or “bridge” for the Austroasiatic populations (or the Tai-Kadai populations) expansion into East Asia [72–75]. Therefore, we collected the first batch of DNA polymorphic profiles of the Hlai language-speaking population utilizing the MPS-based ForenSeq^™^ DNA Signature Prep Kit (DPMA), and multiple population genetic analyses were conducted from different geographic scales incorporation with linguistics contents. It’s extraordinarily notable for clarifying the genetic structures and exploring potential implications for forensic, anthropology and other related subjects.

## 2. Materials and Methods

### 2.1 DNA sample preparation

A total of 136 unrelated healthy Hainan Li (69 females and 67 males) collected from Sanya city and other nine provincial-controlled divisions in south Hainan were sampled and genotyped. The detailed information was presented in Supplementary Table S1 and the specific geographical locations of the samples were shown in Fig.1C. Ethical review for recruitment, sampling and analysis was provided by Hainan Medical University. Written informed consent was provided by all participants, and the inclusion criteria for each volunteer, whose parents and grandparents are aboriginals and have the non-consanguineous marriage of the same ethnical group at least three generations, with the Hlai language as their mother tongue were employed. Peripheral blood samples (PBS) or saliva samples (SS) were collected and dried on an FTA card (Whatman^™^, UK) according to the manufacturer’s protocols.

### 2.2 Library preparation

Libraries were prepared utilizing the ForenSeq^™^ DNA Signature Prep Kit (DAPA) (Illumina^®^, San Diego, CA, USA) in accordance with the recommendations of the manufacturer [76]. For targeted amplification, one punched blood or saliva stain paper (the size was 1.2 mm × 1.2 mm) was used as a PCR template directly for each sample without DNA extraction procedures. The 2800 M control DNA and ddH2O (Promega^®^ Corporation, Madison, WI) were used as positive control and negative control, respectively. All samples were genotyped for DNA profiling with DPMA. Steps for library preparation include target-specific amplification, target enrichment including incorporation of indexed adapters, purification, bead-normalization and pooling, prior to sequencing on the MiSeq Desktop Sequencer using the MiSeq FGx Forensic Genomics System (Illumina^®^, San Diego, CA, USA), all of which were performed following the manufacturer’s recommended protocols [77, 78].

### 2.3 Data processing and genotype calling

Raw data were analyzed using default settings for the Analytical Threshold (AT), Interpretation Threshold (IT), Stutter Filter and Intra-Locus Balance with ForenSeq^™^ Universal Analysis Software (UAS) [79]. Any sequence detected above the analytical threshold of 10 reads was reported to the user for their consideration, while above the 30-read interpretation threshold, the UAS automatically reports the presence of an allele if the overall read depth for the locus was below 650. When ≥ 650 reads were collected for a locus, the AT and IT default analysis parameters were set at 1.5% and 4.5% of the reads for all loci, except for DYS389II (AT = 5.0%, IT = 15%), DYS448 (AT = 3.3%, IT = 10%) and DYS635 (AT = 3.3%, IT = 10%), where noise warranted separate values. Between the AT and IT, the result was flagged by the UAS and the user could determine whether the sequence was an actual variant. The default Stutter Filter percentages for autosomal STR, Y-STR, and X-STR markers ranged from 7.5% (D2S441, D4S2408, PentaD) to 50% (DYS481). User interpretation was also required when the Intra-locus Balance (equivalent to heterozygote balance) felled below 60% for STRs and 50% for iiSNPs, or the level of stutter exceeded the default Stutter Filter value, which varied between STR loci (0-50%). Ambiguous genotypes and allele dropouts were checked by changing the analytical thresholds, in combination with reanalyzing using STRait Razor 3.0 [80] to check bioinformatic concordance of allele calls. The minimal depth of coverage (DoC) was set as 10× for STRs and 5× for SNPs. Alleles of STRs were reported using the nomenclature recommended by Parson et al [81] and the revised Forensic STR Sequence Guide_v3 [82]. The double-check nomenclature were conducted based on our In-house workbooks and manual corrections.

Novel alleles were defined as neither been previously reported in the STR Sequencing Project (STRseq, Accession in NCBI: PRJNA380127) [83] that the purpose of STRseq was to facilitate the description of sequence-based alleles at the Short Tandem Repeat loci targeted in human identification assays nor in previous studies [84–87].

### 2.4 Forensic parameters

#### a) STR/X-STR

Cervus 3.0.7 was employed to calculate the related parameters, including allele frequency, observed heterozygosity (H_obs_), expected heterozygosity (H_exp_), power of discrimination (DP), polymorphism information content (PIC), Hardy-Weinberg equilibrium (HWE), and exclusion probability for duo paternity testing (PE_duo_) and trio paternity testing (PE_trio_), based on both sequence-based and length-based alleles for STR/X-STR profiles [88]. b) **Y-STR:** The Y-STR forensic-associated parameters were conducted as our previous studies [89–91]. Allele and haplotype frequencies were calculated using SAS^®^ 9.4 software. Genetic diversity (GD) was calculated using the following formula by Nei [92]: 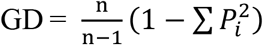, where *n* and *P_i_* denote the total number of the sample and the relative frequency of the *i*-th allele, respectively. Haplotype diversity (HD) was estimated in a similar manner as GD. The discrimination capacity (DC) was defined as the ratio between the number of different haplotypes and the total number of haplotypes. Random match probability (MP) was determined as 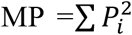, where *P_i_* was the frequency of the haplotype. c) **iiSNP:** STRAF was used to calculate related forensic statistics including: DP, PIC, GD, MP, genotype count (N), power of exclusion (PE), and typical paternity index (TPI) [93]. While, H_obs_, H_exp_, HWE, PE_duo_, and PE_trio_ were conducted by the above-mentioned Cervus 3.0.7 based on iiSNP raw data [88].

### 2.5 Population genetic analyses

We set three different Dataset (I - III) from publicly available population data in literatures and our in-house data for different population analyses from distinct scales. Detailed information of each dataset had been displayed in Table 1.

**Table 1.** The information of three Datasets for different population genetic analyses

| Dataset name | Data | Populations | Population number | individual number |
| --- | --- | --- | --- | --- |
| Dataset I | 92 iiSNPs<br>Raw Data | African ancestry | 7 | 661 |
|  |  | Americas | 4 | 347 |
|  |  | East Asian ancestry | 6 | 640 |
|  |  | European ancestry | 5 | 503 |
|  |  | South Asian ancestry | 5 | 489 |
|  |  |  | 27 | 2640 |
| Dataset II | 17 Y-STRs<br>Raw Data | Sino-Tibetan Family | 29 | 5644 |
|  |  | Sinitic/Chinese | 9 | 2081 |
|  |  | Hmong-Mien Language | 5 | 759 |
|  |  | Tai-Katai/Kra-Dai Language | 9 | 1533 |
|  |  | Tibeto-Burman Language | 6 | 1271 |
|  |  | Austronesian Family | 5 | 1543 |
|  |  | Indo-European Family | 5 | 755 |
|  |  | Austro-Asiatic Family | 1 | 59 |
|  |  | Austro-Asiatic/Sino-Tibetan Family | 2 | 280 |
|  |  |  | 42 | 8281 |
| Dataset III (1) | Raw Data | Y-STR |  | 109142 |
|  |  | Y-STR&Y-SNP |  | 37754 |
| Dataset III (2) | Y-haplogroup | Northern Han | 10 | 558 |
|  | Frequency | Southern Han | 18 | 832 |
|  |  | Hmong-Mien | 23 | 759 |
|  |  | Tai-Kadai | 33 | 1031 |
|  |  | Austronesian | 12 | 251 |
|  |  | Austro-Asiatic | 26 | 930 |
|  |  | Hainan Li | 1 | 476* |
|  |  |  | 123 | 4837 |

\* We selected 489 genotyped Y-STRs of Hainan Li male samples (67 in this study and 422 in our previous study [90]) for the estimation of Y-haplogroups, 476 out of the 489 were observed.

The chordal graph was constructed based on the R Project for Statistical Computing (R, R version 3.5.3) (https://www.r-project.org/) using the “circlize” and “reshape2” packages to show the performances of the DPMA based on the data of the 1000 Genomes Project (1 KG, the data are available at http://grch37.ensembl.org/index.html or https://www.internationalgenome.org/.). The principal component analysis (PCA) was performed with SPSS 22.0 based on the allelic frequencies and raw data of different genetic markers and haplogroup frequencies, and used to explore the extent of correlation of genetic relationships [94]. The Multi-Dimensional Scaling (MDS) was conducted on the basis of allele frequencies by means of R programming language (“stats” and “ggplot2” packages) using different distances (Euclidean and Manhattan). The linear discriminant analysis (LDA) was analyzed by R (“MASS” and “ggplot2” packages) on the basis of 17-YSTR profiles of the Hainan populations. The heatmap was carried by the R Project for Statistical Computing with the package of “ComplexHeatmap” based on allele frequencies and the pairwise fixation index F (*F_st_*) [95, 96], respectively.

To further investigate the genetic structure of populations, a Bayesian model-based clustering approach was adopted with STRUCTURE 2.3.4 [97]. The parameters were set according to the suggestions of Falush et al. [98, 99]. Population structures were inferred by setting the value of the clusters (*K*) from 4 to 7. Ten runs were performed for every *K* value, with an MCMC chain burn-in length of 100,000 iterations followed by 100,000 iterations. The optimum *K* value was identified based on the method proposed by Evanno et al. [100], using the online software STRUCTURE HARVESTER [101]. The STRUCTURE results were plotted using the program DISTRUCT [102].

The *F_st_* [95, 96] and corresponding *P* values between different populations were estimated by analysis of molecular variance (AMOVA) based on the raw data were estimated by Arlequin v3.5 [103]. While, in the calculation of *F_st_* in Dataset II, haplotypes presenting null, intermediate, duplicated or triplicated alleles were removed, and the number of repeats in DYS389I was subtracted from that of DYS389II. Additionally, phylogenetic relationships among different populations were conducted using the Molecular Evolutionary Genetics Analysis 7.0 (MEGA 7.0) software [104] by neighbor-joining (N-J) phylogenetic tree [105] based on the genetic distance matrix (*F_st_* values matrix) [106, 107] and were visualized by the Interactive Tree of Life v4 (iTOL) [108].

### 2.6 Verification with CE-based autosomal STR profiling

All samples were genotyped using The Goldeneye^™^ DNA ID System 20A (Goldeneye^®^ Technology, China), which includes 19 overlapped autosomal STR loci (D19S433, D5S818, D21S11, D18S518, D6S1043, D3S1358, D13S317, D7S820, D16S539, CSF1PO, Penta D, vWA, D8S1179, TPOX, Penta E, TH01, D12S391, D2S1338 and FGA) with the ForenSeq^™^ DNA Signature Prep Kit (DAPA). The experiments were performed according to our previous studies [109, 110] and the manufacturer’s instructions.

## 3. Results and discussion

In this study, 136 individuals from south Hainan island where is the settlements of Li minority were genotyped using the MPS-based ForenSeq^™^ DNA Signature Prep Kit (DAPA), and DNA profiles were obtained from 152 different molecular genetic markers (27 autosomal STRs, 24 Y-STRs, 7 X-STRs, and 94 iiSNPs in total). We evaluated the forensic characteristics and efficiencies of different types of genetic markers included in the ForenSeq^™^ DNA Signature Prep Kit (DAPA) for forensic application in Hainan Li population, and the phylogenetic analyses of the Hlai language-speaking population based on genetics (iiSNPs and Y-STRs), linguistics and other aspects.

### 3.1 Sequencing results

A total of genetic data of 136 Hainan Li samples (69 females and 67 males), generated with ForenSeq^™^ DNA Signature Prep Kit Primer Mix A, were genotyped successfully. As shown in Supplementary Table S2, on the whole, the total Depth of Coverage (DoC) for all genetic markers of 136 samples was 15,941,700 (7,627,759 and 8,313,941 for females and males, respectively; the gender order, similarly hereinafter) and, for each sample, the average DoC of each sample per locus (DoC_s/1_) was 771.17 ± 1363.75 (863.65 ± 1,493.76/F and 816.38 ± 1,239.51/M). The average total DoC for each sample (DoC_s_) was 117,218.38 ± 46,115.29 (110,547.23 ± 39,630.60/F and 124,088.67 ± 51,051.16/M), and the average total DoC for each locus (DoC_1_) was 104,879.61 ± 143,270.46 (59,591.87 ± 91,475.71/F and 54,696.98 ± 70,183.71/M).

For STRs, all the STRs were successfully genotyped, the total success rate of autosomal STRs and the success rate for each STR were 100%. For X-STRs and Y-STRs, the success rates of X-STRs and Y-STRs were 98.00% and 99.19%, except locus DXS10103 and DYS392 (the success rates for the two loci were 89.7% and 85%), the total success rates of X-STRs and Y-STRs reached 99.38% and 99.80%, respectively. For iiSNPs, the total success rate was 92.49% and the success rates for all iiSNPs ranged from 2.94% to 100%, except for rs2269355, rs826472, and rs1736442 (the success rates for the three iiSNP loci were 31.62%, 30.88%, and 2.94%), the total success rate of iiSNPs came up to 94.82%.

As illustrated in Supplementary Table S2 and Fig. 2A-C, the maximum and minimum DoC_l_ for each locus, for STRs, were 6,545.89 ± 3,095.15 for TH01 (6,929.17 ± 2,938.61/F and 6,151.16 ± 3,200.86/M) and 343.07 ± 139.83 for D5S818 (355.45 ± 142.01/F and 332.37 ± 136.72/M), respectively (Fig. 2A(a), Fig. 2B(a), Fig. 2C(a)); for X-STRs, the DoC_l_ ranged from 5,402.13 ± 3,274.00 at DXS7423 (7,367.23 ± 3,207.80/F and 3,378.36 ± 1,757.37/M) to 149.57 ± 107.65 at DXS10103 (211.19 ± 108.66/F and 86.10 ± 58.52/M) (Fig. 2A(b), Fig. 2A(c), Fig. 2B(b), and Fig. 2C(b)), and for Y-STRs from 6,291.19 ± 3,164.73 for DYS438 to 275.06 ± 126.19 for Y_GATA_H4 (Fig. 2A(d), Fig. 2C(c)); for iiSNPs, they spanned from 1,246.40 ± 446.92 for rs1109037 (1,310.75 ± 435.13/F and 1,180.12 ± 449.22/M) to 0.58 ± 3.64 for rs1736442 (0.16 ± 1.31/F and 1.01 ± 4.98/M) (Fig. 2A(e), Fig. 2B(c), and Fig. 2C(d)). It indicated that the DoC_l_ at the same locus had no significant differences no matter in females and males, and the locus imbalances of DoC were observed compared among the same types of loci (in 58 STRs or 94 iiSNPs). Furthermore, the detail DoC_s_ for all samples were presented in Supplementary Table S2 and Supplementary Figure S1-S3, which indicated that the DoC_s_ for all samples were balanceable, with no differences for females and males. Our sequencing results were in accordance with other forensic MPS-related studies [40, 41, 43, 45, 111–115]. No matter in females or males, the DoC_l_ and DoC_s_ had no differences at the same locus, and the DoC were unbalanced in different types of genetic markers.

**Fig. 2.**
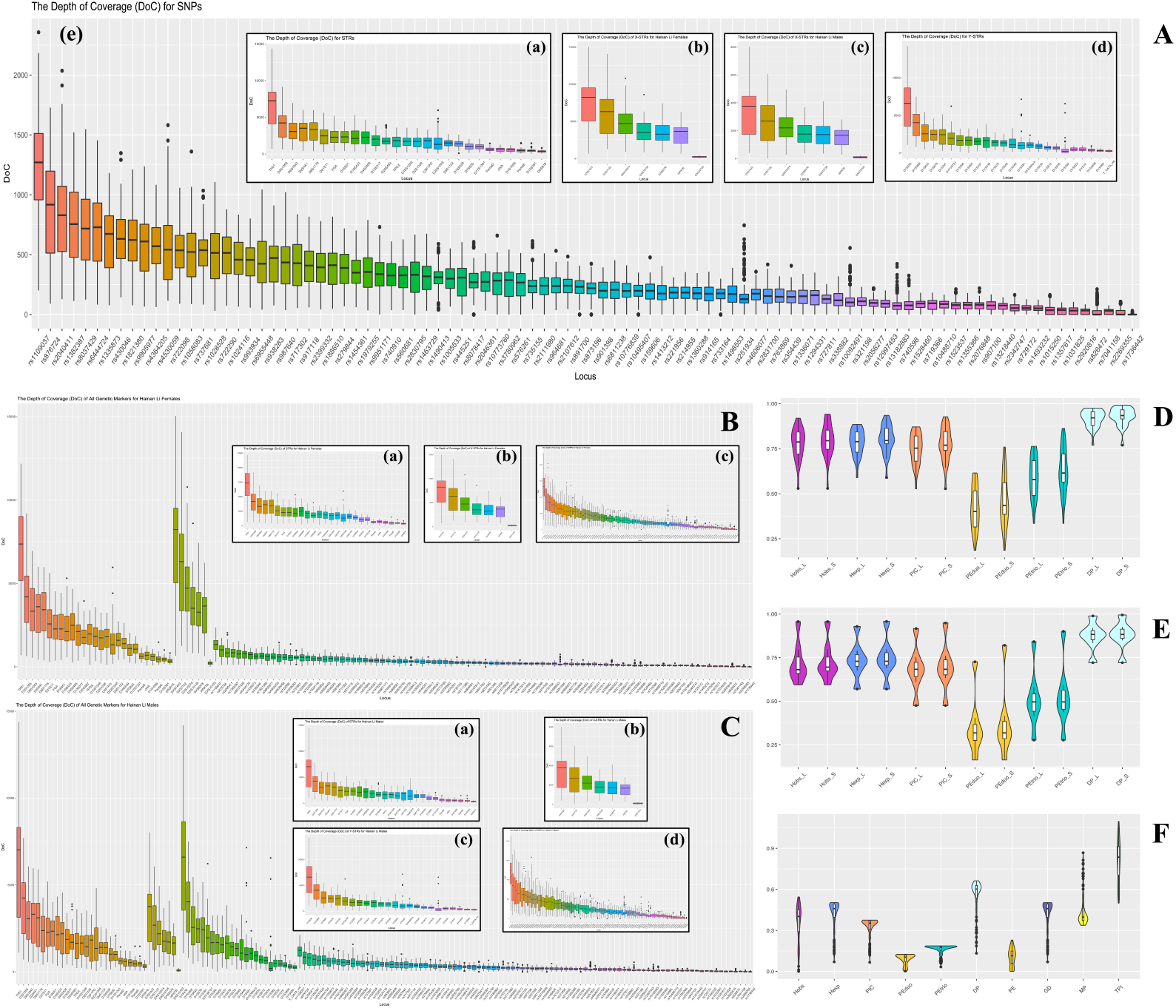
DoC_l_ and forensic parameters of different genetic markers in 136 Hainan Li samples. **A.** Total DoC_l_ in Hainan Li. **(a)** DoC_l_ of 27 autosomal STRs, **(b)** DoC_l_ of 7 X-STRs for females, **(c)** DoC_l_ of 7 X-STRs for males, **(d)** DoC_l_ of 24 Y-STRs, **(e)** DoC_l_ of 94 iiSNPs; **B.** Total DoC_l_ in 69 Hainan Li females. **(a)** DoC_l_ of 27 autosomal STRs in females, **(b)** DoC_l_ of 7 X-STRs in females, **(c)** DoC_l_ of 94 iiSNPs in females; **C.** Total DoC_l_ in 67 Hainan Li males. **(a)** DoC_l_ of 27 autosomal STRs in males, **(b)** DoC_l_ of 7 X-STRs in males, **(c)** DoC_l_ of 24 Y-STRs in males, **(d)** DoC_l_ of 94 iiSNPs in males; **D.** Forensic parameters of 27 autosomal STRs for 136 Hainan Li samples (L: length-based; S: Sequence-based); **E.** Forensic parameters of 7 X-STRs for 69 females (L: lengthbased; S: Sequence-based); **F.** Forensic parameters of 94 iiSNPs for 136 Hainan Li samples.

In addition, we used CE-based method to make the verification for autosomal STRs, the results of 19 autosomal STRs data for 136 samples and 2800M were consistent with the MPS results. Overall, either the quality, the DoC and the amplification positive rates of all genetic markers, or the accuracy of the MPS-based data verified by CE, it was appropriate and acceptable for the sequencing results of Hainan Li in this study.

### 3.2 STR sequence variations observed by the sequence-based method

We made a count for both the sequence-based and length-based alleles in Hainan Li population in Supplementary Table S3-4. Compared with CE methods (length-based genotypes), sequence-based MPS methods could be detected additional STR variations which defined as the alleles with the same length but different sequences for a STR locus, i.e., isoalleles.

The detailed STR sequences and allele frequencies were shown in Supplementary Table S5. Among the 3672 autosomal STR alleles typed in the 136 individuals, 243 distinct length variants and 348 repeat sequence sub-variants were identified across the 27 autosomal STR loci in Hainan Li population. Isoalleles were observed 19 out of 27 autosomal STR loci (70.37%) by MPS-based approach (Supplementary Table S5), and three loci had an obvious allele number increase (increase rate > 100%) for Hainan Li (the increase rates of D12S391, D21S11, and D2S1338 were 227.27%, 210.00%, and 118.18%, respectively) (Supplementary Table S6). According to a previous study in the Torghut and Jalaid Mongol populations [115], the increase rate of D7S820 could reach 114.3% and 166.7% based on the sequencing method, respectively. While, the allele number increase rates of the D13S317 was only 12.5% in Hainan Li population. At D13S317 locus, the allele number increase rate reached as high as 227.27%, however, the allele number increase rates were 107.7% and 141.7% with an obviously differences, respectively. The similar phenomena came under our observation in a series of autosomal STRs, for example, D16S539, D12S391, and D22S1045.

Also, at 7 X-STR loci, we found 82 sequence-based and 57 length-based alleles in females and 57 sequence-based and 49 length-based alleles in males, respectively. In total, four isoalleles were detected revealed by MPS-based method (DXS10135 for both females and males, DXS10074, DXS8378, and DXS10103 only for females). Only DXS10135 locus for females (111.11%) was the obvious allele number increased locus. Under the same basic conditions for populations and sample numbers, the polymorphism, from the sequence diversities in the same length-based STR locus, was significantly higher among females than males.

A total of 156 sequence-based and 119 length-based alleles were identified at the 24 Y-STRs in Hainan Li population. The Y-STR related isoalleles-detected rate of Hainan Li population was 33.33% (8/24), and two loci had an obvious allele number increase for Hainan Li (the increase rates of DYS389II and DYS390 were 160.00% and 100.00%, respectively). While, the increase rates were 100.0% and 240.0% (DYS389II), and 40.0% and 125.0% (DYS390) for Torghut and Jalaid Mongol populations in previous study [115], respectively. Obviously, the same with autosomal STRs, the allele number increase rates for some Y-STR loci in different populations changed without apparent disciplines accordingly.

With the observations of the sequence variations which derived high polymorphisms of some autosomal STR loci in Hainan Li population and other populations that the variations also existed [84, 111–124], it indicated that the sequence variation-derived high polymorphisms widely existed in different populations, and the genetic diversity of the same locus in different populations were diverse under some circumstances. It could be explained by the relatively isolations between populations from the cultural, geographical and social differences, and genetic mutations accumulated over many years in populations.

### 3.3 Forensic parameters for different classes of genetic markers

#### 3.3.1 STR

The forensic parameters for 27 autosomal STR loci based on both sequence-based alleles and length-based alleles in Hainan Li population were shown in Supplementary Tables S7 and Fig. 2D. The number of length-based STR alleles ranged from 5 (TPOX and D4S2408) to 18 (Penta E), and for sequence-based STR numbers from 5 (TPOX) to 36 (D12S391). The H_exp_ for length-based STRs in Hainan Li spanned from 0.5890 at TPOX to 0.8850 at D18S51, and from 0.5890 at TPOX to 0.9350 at D12S391 for sequence-based STRs. The PIC ranged from 0.5270 (TPOX) to 0.8710, and from 0.5270 (TPOX) to 0.9270 for length-based and sequence-based STRs, respectively. The top three most polymorphic loci were D18S51(0.871), Penta E (0.861), and D2S1338 (0.854) for length-based STRs, for sequence-based loci, the most top three most polymorphic loci were D12S391 (0.9270), D21S11 (0.9050), and D2S1338 (0.8910). All DPs were greater than 0.7690 (TPOX), and the highest DP values were detected at D18S51 (0.975 for length-based STR) and D12S391 (0.9910 for sequence-based STR).

#### 3.3.2 X-STR

Moreover, for X-STRs in females (presented in Supplementary Tables S7 and Fig. 2E), the H_exp_ values were in a range of 0.5700 (DXS7423) to 0.9290 (DXS10135), and 0.5700 (DXS7423) to 0.9580 (DXS10135) for length-based and sequence-based STR loci, respectively. The values of PIC varied from 0.4750 (DXS7423) to 0.9170 (DXS10135) for the length-based X-STR, and from 0.4750 (DXS7423) to 0.9490 (DXS10135) for the sequence-based X-STRs. Additionally, the minimum value of DP was 0.7210 at the same DXS7423 locus, whereas the maximum DP values were 0.9890 and 0.9950 at DXS10135 for length-based and sequence-based X-STR loci, respectively.

#### 3.3.3 Y-STR

As for Y-STRs, the length-based and sequence-based Y-STR haplotype profiles and related forensic parameters were presented in Supplementary Tables S8. The GDs, which had differences in 6 out of 24 Y-STRs (25%), were 0.8485 and 0.6984 (DYS389II), 0.7888 and 0.6486 (DYS390), 0.8218 and 0.7752 (DYS612), 0.9502 and 0.9055 (DYF387S1), 0.8078 and 0.7761 (DYS481), and 0.7843 and 0.7684 (DYS635) for sequence-based and length-based Y-STR loci, respectively. The DC, MP, and HD, with no differences between sequence-based and length-based Y-STR loci, were 0.9851, 0.0153, and 0.9998, respectively.

#### 3.3.4 iiSNPs

The genotypes, allele frequencies and related forensic parameters of 94 iiSNPs were shown in Supplementary Tables S9-10 and Fig. 2F. The Minor Allele Frequency (MAF, frequency of the second most frequent allele) ranged from 0.0357 (rs826472; T) to 0.4963 (rs1109037, G; rs987640, A). We found eight SNP loci with unbalance values (the minor locus allele frequency was blow 0.1) in Hainan Li, they were respectively rs826472 (T; 0.0357), rs251934 (G; 0.0682), rs740910 (G; 0.0735), rs729172 (T; 0.0796), rs735155 (G; 0.0846), rs1024116 (T; 0.0882), rs7041158 (T; 0.0962), and rs737681 (T; 0.0993). The H_exp_ for 94 iiSNPs ranged from 0.0700 (rs826472) to 0.5030 (rs1015250) with an average H_exp_ for 0.4146 (± 0.1108). The minimum PIC was 0.0665 at rs826472, and the maximum PIC was 0.0750 at rs1015250, rs354439, rs987640 and rs1109037, while, the average PIC was 0.3212 ± 0.0741. The average DP was 0.5494 (± 0.1263) with a range DP of 0.1327 (rs826472) to 0.6626 (rs3780962). The PE varied from 0 (rs1736442) to 0.2291 (rs1463729), and the average PE value was 0.1063 (± 0.0627). For GD, the range was 0.0697 at rs826472 to 0.5029 at rs1015250 (0.4146 ± 0.1107). The maximum and minimum of MP values were rs826472 (0.8673) and rs3780962 (0.3374), respectively. The TPI spanned 0.5000 (rs1736442) to 1.0968 (rs1463729) with an average of 0.8079 ± 0.1545.

In total, as shown in Table 2, based on sequence-based and length-based alleles, the cumulative discrimination power (CDP) of the 27 autosomal STRs were 1-1.44 × 10^−34^ and 1-8.98 × 10^−32^, and the CDP of the 7 X-STR were 1-2.33 × 10^−08^ and 1-6.80 × 10^−08^, respectively. The combined matching probability (CMP) for sequence-based and length-based alleles were, 1.39 × 10^−34^ and 9.07 × 10^−32^ for the 27 autosomal STRs, 2 × 10^−8^ and 7 × 10^−8^ for the 7 X-STR loci. For iiSNPs, the CDP and CMP were 1-1.80 × 10^−34^ and 1.81 × 10^−34^, respectively. In addition, for the length-based 27 autosomal STRs, the combined power of exclusion (CPE) were 1-2.37 × 10^−07^ and 1-1.28 × 10^−11^ for duo and trio paternity testing, respectively, in Hainan Li; correspondingly, the CPE of duo and trio paternity testing for sequence-based 27 autosomal STRs were 1-1.87 × 10^−08^ and 1-5.05 × 10^−13^, respectively. While, the CPE of the 94 iiSNPs were 1-1.17 × 10^−04^ and 1-6.57 × 10^−08^ for duo and trio paternity testing, respectively. Thus, corresponding to an increase in the CDP, a decrease in the CMP was observed on account of the increasing diversity of sequence-based alleles. Hence, the 27 autosomal STRs (no matter sequence-based or length-based STR loci) and 94 iiSNPs included in ForenSeq^™^ DNA Signature Prep Kit (DAPA) provided high polymorphism information content and could be used to perform forensic individual identification and parentage testing in Hainan Li population separately or federatively. Furthermore, even though there were no significantly differences for CDP and CMP for the sequence-based 27 autosomal STRs and 94 iiSNPs, the CPE of 27 sequence-based autosomal STRs for duo and trio paternity testing were higher than the set of 94 iiSNPs in Hainan Li population, which indicated that the efficiencies for the set of sequence-based 27 autosomal STRs were better than the set of 94 iiSNPs in parentage testing.

**Table 2.** Forensic parameters for autosomal STRs, X-STRs for females, and iiSNPs

|  | Autosomal STR |  | X-STR for females |  | iiSNP |
| --- | --- | --- | --- | --- | --- |
|  | Length-based | Sequence-based | Length-based | Sequence-based |  |
| Number of markers | 27 | 27 | 7 | 7 | 94 |
| Cumulative DP (CDP) | $1-8.98 \times 10^{-32}$ | $1-1.44 \times 10^{-34}$ | $1-6.80 \times 10^{-08}$ | $1-2.33 \times 10^{-08}$ | $1-1.80 \times 10^{-34}$ |
| Cumulative MP (CMP) | $9.07 \times 10^{-32}$ | $1.39 \times 10^{-34}$ | $7.00 \times 10^{-08}$ | $2.00 \times 10^{-08}$ | $1.81 \times 10^{-34}$ |
| Cumulative PE<sub>duo</sub> (CPE<sub>duo</sub>) | $1-2.37 \times 10^{-07}$ | $1-1.87 \times 10^{-08}$ | $1-3.28 \times 10^{-02}$ | $1-1.97 \times 10^{-02}$ | $1-1.17 \times 10^{-04}$ |
| Cumulative PE<sub>trio</sub> (CPE<sub>trio</sub>) | $1-1.28 \times 10^{-11}$ | $1-5.05 \times 10^{-13}$ | $1-3.71 \times 10^{-03}$ | $1-2.04 \times 10^{-03}$ | $1-6.57 \times 10^{-08}$ |

### 3.4 Genetic differentiation based on iiSNPs

In this study, we set up three Datasets including iiSNPs, Y-STR, and Y-haplogroup data of diverse populations with different ancestries and language families to perform population genetic analyses separately from different perspectives. As shown in Dataset I (Table 1), a total of 27 populations, twenty-six populations with five different ancestries from the 1000 Genomes Project [125–128] and Hainan Li population, included 2640 individuals (2504 individuals for 1 KG populations and 136 individuals for Hainan Li) with raw data of 94 iiSNPs to make population comparisons. (Detailed geographic and ancestry information were presented in Fig. 1A and Supplementary Table S11.)

The chordal graph in Fig. 3A and Supplementary Table S12 showed the basic information and performances of 94 iiSNP from the ForenSeq^™^ DNA Signature Prep Kit (DAPA) in 1 KG populations. The set of iiSNPs spread over 22 autosomal chromosomes with a range of 2 (chromosome 19) to 6 (chromosome 1) iiSNPs for each, and the most severe consequences of this iiSNPs were intergenic variant (47), intron variant (34), regulatory region variant (8), non-coding transcript exon variant (3), 3 prime UTR variant (2), and TF binding site (1). (for rs740598, it located in regulatory region variant/intergenic variant) Moreover, the percent of the ancestral alleles for the set of 94 iiSNPs were 35.10% for C (33), 24.45% for G (23), and 20.21% for A (19) and T (19), respectively. In addition, the Minor Allele Frequency (MAF, frequency of the second most frequent allele in 1000 Genomes Phase 3 combined population) varied from 0.24 (A) for rs10495407 to 0.50 for four iiSNPs (rs1490413, rs3780962, rs722290, and rs722098), with an average MAF for 0.3851 ± 0.0789 in the set of 94 iiSNPs, and the highest population MAFs for the 94 iiSNPs were in the range of 0.3 at rs2056277 to 0.5 at 37 iiSNPs. No matter the equilibrium of distribution or the stability of hereditary, additionally taking the results of allele frequencies and forensic parameters (CDP, CMP and CPE) into considerations, the efficiency of the set of 94 iiSNPs selected in ForenSeq^™^ DNA Signature Prep Kit performed relatively well in forensic application scenarios. Thus, except for rs1736442 (The amplification positive rate was only 2.94% in Hainan Li) and rs938283 (The related information didn’t release for 26 populations in 1000 Genomes Phase 3), a total of 92 iiSNPs were recruited for population genetic analyses between Hainan Li and 1 KG populations.

**Fig. 3.**
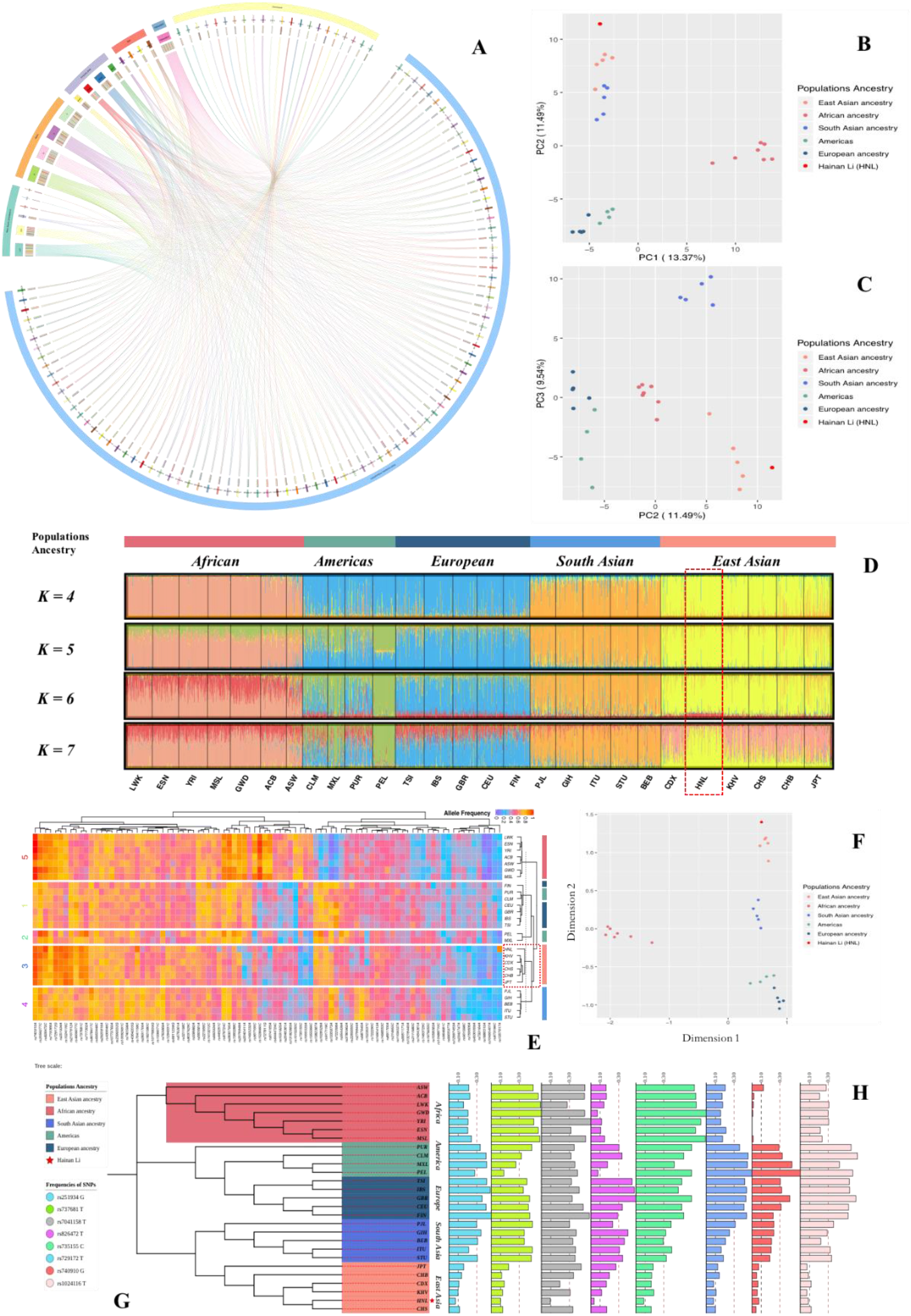
Population genetic analyses based on iiSNPs in Dataset I. **A.** Chordal graph of 94 iiSNPs (Lo1, intergenic variant; Lo2, intron variant; Lo3, regulatory region variant; Lo4, non-coding transcript exon variant; Lo5, TF binding site; Lo6, 3 prime UTR variant); **B.** PCA (PC1 and PC2); **C.** PCA (PC2 and PC3); **D.** Structure analysis (Hainan Li in red dotted box); **E.** Heatmap (HNL, Hainan Li; EAS and HNL in red dotted box); **F.** MDS based on Euclidean distance; **G.** N-J tree (The red star represented Hainan Li); **H.** Unbalanced loci of Hainan Li (MAF <0.10).

#### 3.4.1 Dimensionality deduction analyses

Dimensionality reduction analyses, including PCA, LDA, MDS, LE (Laplacian Eigenmaps) and LLE (Locally linear embedding), can accelerate the speed of algorithm execution, improve the performance of analysis model and reduce the complexity of data at the same time. To illustrate the genetic landscapes of populations in Dataset I (Table 2), especially for Hainan Li population, the dimensionality reduction analyses (PCA and MDS) were conducted based on the frequencies of 92 iiSNPs which were illustrated in Supplementary Table S13.

The PCA provides a method of visualizing the essential patterns of genetic relationships, and allows us to identify and plot the major patterns within a multivariate data set in order to indicate that the populations with closer geographical distances have more intimate relationships. As shown in Fig. 3B and Fig. 3C, the first, second, and third components (PC1, PC2, and PC3) accounted for 13.37%, 11.49%, and 9.54% of the total variance observed within these populations, respectively. In the PCA diagram, populations from five different intercontinental ancestries clustered separately, East Asian and South Asian ancestry populations clustered together on the upper left and the two groups separated relatively, populations from Americas and European ancestry got together on the left bottom, and the Africa-originated populations clustered together, distributed on the middle right isolated. The PCA analysis made a relatively clear distinction between groups of intercontinental ancestries, and Hainan Li population distributed in a relatively isolated location, at the extreme top left corner, attached to East Asian ancestry populations.

While, in order to make further confirmation about the relations between Hainan Li and populations in 1 KG conducted by PCA in Fig. 3B-C, the MDS, with Euclidean distance, was conducted in Fig. 3F, which depicted the genetic relationships between Hainan Li and the 1 KG populations. Similar to the PCA in Fig. 3B, Hainan Li located at the upper right corner (at the upper left corner in PCA) in MDS diagram, cluster with EAS ancestry populations at intervals.

The dimensionality reduction analyses (PCA and MDS) conducted based on the frequencies of 92 iiSNPs in ForenSeq^™^ DNA Signature Prep Kit illustrated that Hainan Li had an East Asian ancestry and had relatively far distances with other EAS populations (1 KG) in the inner EAS cluster, indicating Hainan Li was an isolated population relatively compared with 1 KG populations. The geographic isolation and the indigenous cultures protection led Li population to inhabit and live in south Hainan island which separated by Qiongzhou Strait and Beibu Gulf apart from Chinese mainland and Southeast Asia mainland during the Last Glacial Maximum (~14.5kya) [6–8, 27, 28], resulting in limited gene flows with populations in Hainan island and little gene flows with the populations outside Hainan island.

#### 3.4.2 STRUCTURE analysis

STRUCTURE analysis is commonly recognized to be capable of inferring population structure and assigning individuals to populations using multi-locus genotypic data [97]. In present study, STRUCTURE clustering analysis was performed to reflect the memberships of biogeographic ancestry components for Hainan Li and 1 KG populations with the number of hypothetic populations (*K*) defined at 4-7 based on the frequencies of 92 iiSNPs (Shown in Supplementary Table S13). And a burn-in period of 100,000 was also taken into account to acquire representative estimations of the parameters.

The results for *K* = 4 to 7 were shown in Fig. 3D, which population names as well as their corresponding population ancestries were labeled on the bottom and the top of the figure, and the width of each bar was proportional with the population sample size, we observed a plateau of the estimated posterior probability at *K* = 5. According to the guidelines in STRUCTURE manual, *K* = 5 was the most appropriate value for the STRUCTURE analysis between Hainan Li and populations in 1 KG. At *K* = 5, the five distinct ancestry groups could be distinguished from each other. For AMR and EUR populations, though the genetic differences were obvious to some extent, the genetic relationships were ambiguous from the STRUCTURE plot, especially for PUR and CLM from AMR when compared with EUR populations. While, populations from other ancestries (AFR, SAS, and EAS) had marked difference, and the accurate biogeographic ancestry relationships revealed by STRUCTURE were consistent with the PCA and MDS results. For the inner of the EAS, the components of Hainan Li have significant differences with other EAS populations (When *K* = 6 and 7, especially for 7). At *K* = 7 for EAS populations, the components of Hainan Li had significant variance with other EAS populations. It indicated that the set of 92 iiSNPs had good ability to distinguish populations from diverse intercontinental groups (merely for intercontinental groups), however, it has no ability to distinguish the inner-populations in the same continental scale. In addition, Hainan Li was a population with East Asian ancestry, with their own characteristic genetic features to some extent.

#### 3.4.3 Heatmap

Heat map, with abundant color changes and vivid information expressions, is widely used in various big data analysis scenarios. The heat map visually shows the density or frequency of different types of data. Based on the frequencies of 92 iiSNPs which were presented in Supplementary Table S13, a heat map was contrasted to demonstrate the dendrogram relationships between Hainan Li and populations from 1 KG in Fig. 3E. As a whole, all populations of Dataset I were clustered into two large clusters (AFR and others). The clustering relationships could provide circumstantial evidences for the out-of-Africa hypothesis (Eve hypothesis) [129–141]. Except the AFR cluster, the other cluster could separate two main sub-clusters (European and Asian cluster). For European sub-cluster, EUR and AMR populations got together without distinct distinction. The PEL and MXL from AMR clustered in a separated branch, and the PUR and CLM with the same AMR ancestry clustered together with other EUR ancestry populations. With the discovery of America by Christopher Columbus (1451-1506) in the year of 1492 [142, 143], large numbers of Europeans swarmed into the continent of America for lives and expansions, and immigrated, settled, thrived, and prospered from the beginning to now [144–146]. The exact course of history manifested the comprehensive and deep gene flows existed between the immigrants and the aborigines in America continent which appropriately explain the clustering of both in Heatmap. While, for Asian sub-cluster, EAS and SAS populations located in two different branches, respectively. In the red dotted box of Fig. 3E, Hainan Li, similar with JPT, isolated in an insular branch separately and clustered with other EAS populations, which indicated that Hainan Li was an isolated population in East Asia.

#### 3.4.5 Phylogenetic analyses

The *F_st_* and the corresponding *P* values between every two populations in Dataset I were calculated in Supplementary Table S14. The two maximum *F_st_* values observed between ESN and PEL (0.1214), and MSL and PEL (0.1217), correspondingly, the two minimum values were 0.0007 between TSI and IBS, and 0.0011 between CHS and CHB. As for population differentiation based on *F_st_* values, the genetic distance increased relative to the magnitude of geographical separation in most situations. Hainan Li showed the smallest differentiation with the KHV (*F_st_* = 0.0015, *P* = 0.03809 ± 0.0056) and CHS (*F_st_* = 0.0026, *P* = 0.01074 ± 0.0033). Furthermore, the MXL population (*F_st_* = 0.0197, *P* < 0.0001) had the most distant relationship with the Hainan Li. Except for CHS and KHV, significant genetic differences were observed between the Hainan Li and the populations from the other intercontinental populations (all *P* values were less than 0.05).

To further evaluate the genetic structure relationships among these populations, we constructed the phylogenic tree using the N-J method based on genetic distances in Fig. 3G. The EAS, SAS, and AFR ancestry populations clustered in different branches respectively, while EUR and AMR clustered in same branch and relatively separated in the inner. The results were in line with the clustering relationships illustrated by Heatmap, on account of the all-around and underlying gene flows between both because of the expansion of Europeans into the continent of America. What’s more, for the East Asian ancestry populations, the JPT population was selected as an outgroup. The remaining EAS populations and Hainan Li population clustered together, indicating their homogeneous origin. For details, CDX and KHV (*F_st_* = 0.0020) got together, and CHS and HNL clustered together, they all belong to Southeast Asia from the perspective of geography. While, CHB has a relatively far distances with other populations. From the respect of linguistics and language families, Chinese (CHS and CHB), Li and Dai language (Tai-Kadai/Kra-Dai Language for HNL and CDX) belong to the Sino-Tibetan Language Family, and the classification of Gin language (KHV) is not totally confirmed between Sino-Tibetan Language Family and Austro-Asiatic Language Family. The genetic distances indicated by *F_st_* values and the phylogenic relationships based on N-J tree were consistent with the results of above-mentioned genetic analyses (PCA, MDS, and cluster analysis). Therefore, the genetic distances were consistent with geographic scales in this study to some degree.

As a result, from the perspective of the linguistics or genetics, Hainan Li (Hlai, a branch of Tai-Kadai Language which belongs to Sino-Tibetan Language Family) had a relatively close relationship with the KHV (Gin language, Sino-Tibetan/Austro-Asiatic Language Family, *F_st_* = 0.0015, *P* = 0.03809 ± 0.0056) and CHS (Chinese, Sino-Tibetan Language Family, *F_st_* = 0.0026, *P* = 0.01074 ± 0.0033).

In conclusion, from the genetic analyses based on 92 iiSNPs of 2640 individuals from five different intercontinental ancestries, for some diverse intercontinental populations, the set of 92 iiSNPs (especially for eight SNPs presented in Fig. 3H, with apparently disequilibrium in Dataset I populations and the MAF below 0.10 in Hainan Li) also could be applied to the estimation of the biogeography ancestry from the intercontinental scale, which extended the forensic applications of the ForenSeq^™^ DNA Signature Prep Kit in Hainan Li and 1 KG populations, and the Tai-Kadai language-speaking Hainan Li, with East Asian ancestry and their own characteristics in genetics, was an relative isolated population and had a relatively close relationship with the KHV and CHS.

### 3.5 Population differentiation based on Y-STRs

Furthermore, to make additionally population differentiations between Hainan Li and populations which had relatively close distances in geographical scale, and populations in Hainan island [89, 91], we collected publicly published Y-STR data in literatures and our house-in Hainan Y-STR dataset for Dataset II (in Table 1) to evaluate the differences between Hainan populations including Hainan Li and the populations from south China [89, 91, 147–158], Southeast Asia [159–162] and Australia [163] with different language families. The details of Dataset II, a total of 8281 individuals with 17 Yfiler data (DYS19, DYS389I, DYS389II, DYS390, DYS391, DYS392, DYS393, DYS385a/b, DYS437, DYS438, DYS439, DYS448, DYS456, DYS458, DYS635 and Y_GATA_H4), were presented in Supplementary Table S16 and the elaborate geographic positions of the 42 populations which belonged to five language families were orientated in Fig. 1B.

#### 3.5.1 Principal component analysis

In Dataset II, most populations came from different administrative divisions of China in the south and the countries in southeast Asia, which had a relatively close geographical distances with Hainan island. The PCA based on all raw data of 8281 individuals (Supplementary Table S16) was presented in Supplementary Figure S4, indicating that the mixture phenomenon at the relatively narrow and local areas in Dataset II was existed. Moreover, in order to make clearer relationships among Dataset II populations, the PCA based on the allele frequencies of 17 Yfiler, the PCA were performed and shown in Fig. 5A and Fig. 5B, the first three components (27.04% totally) which accounted the proportions of the total variances observed within these populations in Dataset II were 10.05%, 8.77%, and 8.22%, respectively. In general, the intimate and complicated genetic differentiations among populations were also existed, while, the populations with the same language family aggregated in PCA plot to some extent. Further, from a relatively smaller scale and perspective, Hainan Han had a relatively close distance with other Han Chinese populations, and Hainan Lingao clustered with populations which belongs to Tai-Kadai language families (Fig. 5A). What’s more, on the basis of different principal components, Hainan Li located in a relatively isolated situation in PCA diagrams (both in Fig. 5A and Fig. 5B). So as to estimate the internal population relationships and structures precisely, we utilized the 17 Yfiler raw genotype data, taking each individual into account, to evaluate genetic differences from different scales.

First of all, the populations in Dataset II were divided into three groups: 1) Group I: according to the administrative divisions in south China, including Guangdong (2 populations), Guangxi (7), Guizhou (4), Hunan (4), Sichuan (3), and Yunnan (4); 2) Group II: the populations from the same nationality, containing Han (7 populations from different provinces), Hui (2), Miao (3), Yao (2), Yi (3), and Gelao (2); 3) Group III: the populations from other countries of southeast Asia and Australia, consisting of Australia (3), Malaysia (3), Sigapore (3), and Vietnam (3). Then, the principal component analyses were performed for different groups, and the PCA results were presented in Fig. 4A-F for Group I, Fig. 4G-L for Group II, and Fig. 4M-P for Group III, respectively. It demonstrated that, from the perspectives of geographic and ethnic scales, different nationalities and diverse administrative divisions of south China and other countries, the local populations couldn’t divide into isolated parts relatively and apparently. However, for the isolated Hainan island, the populations were relatively isolated on the PCA plot (Fig. 4Q), especially for Hainan Li.

**Fig. 4.**
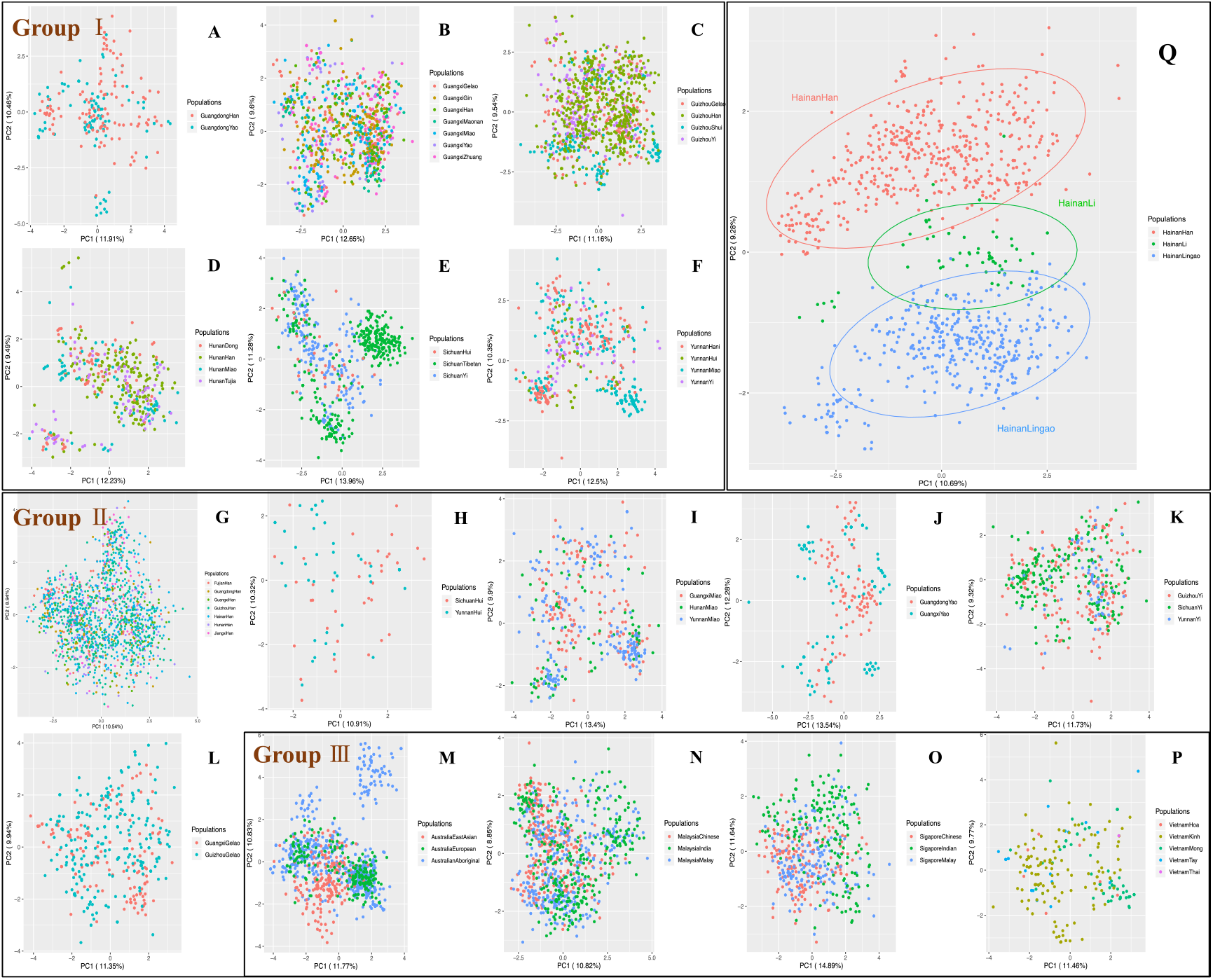
PCA diagrams of different Groups based on 17 Yfiler raw data. **Group I: A.** Guangdong (Han and Yao); **B.** Guangxi (Han, Gelao, Gin, Maonan, Yao, and Zhuang); **C.** Guizhou (Han, Gelao, Shui, and Yi); **D.** Hunan (Han, Dong, Miao, and Tujia); **E.** Sichuan (Hui, Tibetan, and Yi); **F.** Yunnan (Hani, Hui, Miao, and Yi); **Group II: G.** Han Chinese (Fujian, Guangdong, Guangxi, Guizhou, Hainan, Hunan, and Jiangxi); **H.** Hui (Sichuan and Yunnan); **I.** Miao (Guangxi, Hunan, and Yunnan); **J.** Yao (Guangdong and Guangxi); **K.** Yi (Guizhou, Sichuan, and Yunnan); **L.** Gelao (Guangxi and Guizhou); **Group III: M.** Australia (East Asian, European, and Aboriginal ancestries); **N.** Malaysia (Chinese, Indian, and Malay ancestries); **O.** Sigapore (Chinese, Indian, and Malay ancestries); **P.** Vietnam (Kinh, Mong, and Toy; Thai and Hoa which were presented in PCA plot didn’t bring into Dataset II due to the sample sizes.); **Q.** Hainan (Li, Han, and Lingao).

Due to three massive waves of north-to-south migrations and other smaller southward migrations in Chinese history, the expansions of Han Chinese accelerated the communications and gene flows with the southern natives, including those speaking Tai-Kadai, Austro-Asiatic and Hmong-Mien languages [29, 30, 164, 165]. The geographical isolation by the formation of the Qiongzhou Strait during the period of the LGM (~14.5kya) [4, 8, 21, 27, 28] resulted in the limited gene flows between Hainan populations and the surrounding populations in East and Southeast Asia, and the culture isolation led to limited exchanges of genes and cultures between Hainan Li and other Hainan groups in the population evolutionary and adaptation process.

#### 3.5.2 Multi-dimensional scaling

In this study, the MDS plots were conducted based on the allele frequencies by Euclidean distance and Manhattan distance respectively, which characterized the genetic relationships between Hainan Li population and other forty-one populations mentioned in Dataset II. As shown in Fig. 5C-D, each population was represented by a small dot with different color in the multidimensional space, and the distances between the small dots showed the genetic relationships among the populations with distinct language families. The results of MDS analysis by Euclidean distance (Fig. 5C) had no apparently differences with the principal component analysis in Fig. 4A. Moreover, in the Manhattan distance-based MDS plot (Fig. 5D), Hainan Lingao was intertwined with some Sinitic/Chinese and Tai-Kadai language-speaking populations. On the contrary, Hainan Han clustered with Guizhou Han located in a relatively isolated place. In addition, Hainan Li in Manhattan MDS plot isolated with other populations were in accordance with the performances in the MDS conducted by Euclidean distance and PCA plots which presented in Fig. 5D, 5A, and 5B. What’s more, we collected all raw data of 457 Hainan Han, 450 Hainan Lingao, and 67 Hainan Li to make MDS analyses with Euclidean distance and Manhattan distance (Supplementary Figure S5. A-B), the results illustrated the isolated relationships among Hainan populations.

**Fig. 5.**
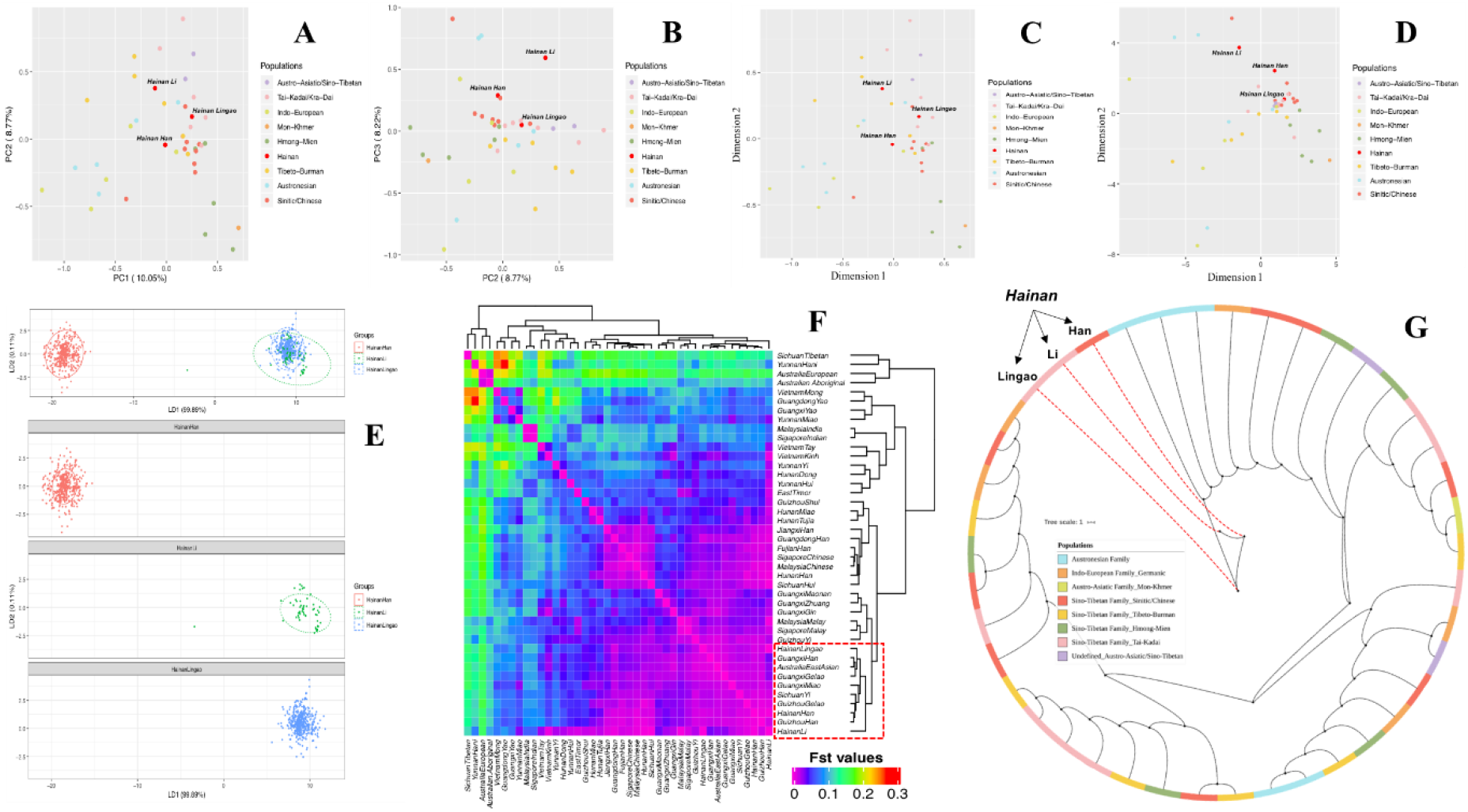
Population genetic analyses based on Y-STRs in Dataset II. **A.** PCA (PC1 and PC2); **B.** PCA (PC2 and PC3); **C.** MDS based on Euclidean distance; **D.** MDS based on Manhattan distance; **E.** LDA of Hainan populations; **F.** Heatmap based on *F_st_* values matrix; **G.** N-J phylogenetic tree.

#### 3.5.3 Linear discriminant analysis

From the above PCA and MDS analyses, the populations in Hainan island showed the relatively separate relations for each other. The Y-STR raw data of Hainan populations were conducted to implement the data dimensionality reduction method with supervise, and obviously, as shown in Fig. 5E, Hainan Li and Lingao (belongs to Tai-Kadai language) clustered together, and Hainan Han was separated by the Tai-Kadai cluster. Separately views for the LDA results, the inner connections for each Hainan populations were tightly. Differences from the results presented by PCA and MDS were that Hainan Li and Lingao had an intimate relationship in LDA. Linguistically speaking, Hainan Han used the Hainanese/Chinese (belongs to Sinitic Language) as daily communications, both the Hlai language used by Hainan Li and the Ong Be language used by Lingao belongs to Tai-Kadai language, moreover, all three belongs to Sino-Tibetan Language Family. Therefore, the relationships manifested by LDA between Hainan populations, especially for Hainan Li and Lingao, could be explained from the perspective of anthropological linguistics reasonably.

#### 3.5.4 *F_st_* and phylogenetic analyses

The detail information of calculations of pairwise *F_st_* and the corresponding *P* values between the Hainan Li and other 41 populations from different administrative divisions of south China and Southeast Asia in Dataset II on the basis of Yfiler haplotypes were indicated in Supplementary Table S16. There were significantly genetic differences between Hainan populations and other populations in Dataset II. (*P* < 0.05) While, Hainan Li had no differences with Hainan Lingao (*F_st_* = 0.0011, *P* = 0.99902 ±0.0002) and Hainan Han (*F_st_* = 0.0028, *P* = 0.99902 ± 0.0002). Based on the symmetric *F_st_* values matrix, a heat map was conducted to measure the genetic differentiations among the populations in Dataset II presented in Fig. 5F. The color of each block deepened with the corresponding *F_st_* value. The color scale ranged from deep pink for the lowest *F_st_* value to red for the highest *F_st_* value. It clearly indicated that Hainan populations clustered together in a small bunch (in the red dotted rectangle). Hainan Han and Guizhou Han clustered together and Hainan Lingao relatively isolated in another sub-cluster, while, Hainan Li secluded at an isolated cluster independently in the same punch. The relationships based on the clustering analysis were in line with the results of PCA and MDS.

Furthermore, the phylogenetic relationships among populations from Hainan populations and other populations in Dataset II were illustrated by an N-J phylogenetic tree. Fig. 5G illustrated the phylogenetic relationships between Hainan populations and other populations in Dataset II, and the details of the phylogenetic N-J tree were shown in Supplementary Figure S6. Different colors represented distinct populations with diverse language families. In general, the structures uncovered by phylogenetic analysis obviously demonstrated Hainan populations isolated in different adjacent branches, and clustered together relatively, and Hainan Li had a closer distance with Hainan Lingao compared with Hainan Han. What’s more, populations in East and Southeast Asia couldn’t separate and cluster by language families absolutely. The population expansions and mixtures led to the interracial marriages and gene flows, which generated the complicated genetic structures in East and Southeast Asia.

From the respect of linguistics and language families, the Hlai language (Li, Hainan) [31, 32] and Ong Be language (Lingao, Hainan) [166–168] are the different categories of Tai-Kadai language [169–174], in this language branch, another important category is Sinitic/Chinese language, which including varieties of Han Chinese languages (including Hainanese) [175–177]. Hlai, Ong Be, as well as Hainanese all belongs to Sino-Tibetan Language Family from the classification of language perspective. By and large, the same language used by different populations always has a directivity to the same and similar ancestry population, which meant that genetic ancestry is strongly correlated with linguistic affiliations as well as geography [74]. No matter from the perspective of genetics or linguistics, as an isolated population in isolated Hainan island relatively, Hainan Li had a close relationship with Hainan Lingao, and a relatively far distance with Hainan Han.

### 3.6 The prediction of Y-haplogroups

It is generally known that no algorithms could be powerful enough to distinguish the same or very similar haplotypes and assign them into different haplogroups and the convergence of Y chromosome STR haplotypes among different haplogroups has compromised the accuracy of haplogroup prediction. Thus, we used our in-house Dataset III (1), including 37,754 pieces of Y SNP/STR data and 109,142 Y-STR in total which mainly from East and Southeast Asia, to make more precisely predictions for 67 Hainan Li males in this study plus the 422 Y-STR profiles which had been reported by our previous study [90, 178]. We selected the Y-STR data of 67 male samples in this study and 422 males in our previous study [90] for the estimation of Y-haplogroups of Hainan Li populations. Eventually, 476 out of the 489 genotyped Y-STRs (97.34%) were observed in the derived state which were presented in Supplementary Table S18, thus defining 6 haplogroups observed in our samples, belonging to major clades O1 and O2. The predominant haplogroups were O1b1a1a-M95 (45.40%), O1a-M119 (26.58%), O1b1a1a1a1a1-M88 (17.38%), O2-M122 (4.09%), O2a1b-IMS-JST002611 (2.01%), and O2a2b1a1-M117 (1.84%). (determined according to ISOGG 2019, https://isogg.org/tree/) Our results were in accordance with previous studies [12, 16], M95 was the predominant haplogroup with a range from 46% to 91.94% for the proportions in different branches of Hainan Li.

To discern the detailed relationship between the Hlai language-speaking population and other related populations with diverse language families, we established the Dataset III (2) which was presented in Table 1 to conduct the bioinformatics analysis. A PCA graph was performed among 123 populations of Dataset III (2) including Tai-Kadai, Hmong-Mien, Austro-Asiatic, Austronesian, and Chinese (Southern and Northern Han) populations from East and Southeast Asia, 4837 individuals in total [165, 179–182]. The first three principal components could explain 29.87% of the total variances. (The PC1, PC2, and PC3 accounted for 12.25%, 9.54%, and 8.08%, respectively.) From the dendrogram of Fig. 6A, the populations with different language branches were separated relatively, and Hainan Li had a close relationship with Tai-Kadai cluster. While in Fig. 6B, the situation got complicated and the populations with different language families mingled with each other, especially for Tai-Kadai and South Han with no apparently difference, Hainan Li was exactly located in this region. The interpopulation comparison demonstrated that Hainan Li had close affinity with Tai-Kadai language-speaking populations based on the evidences of Y-haplogroups which contained information about the subsequent colonization, differentiations, and migrations overlaid on recent population ranges.

**Fig. 6.**
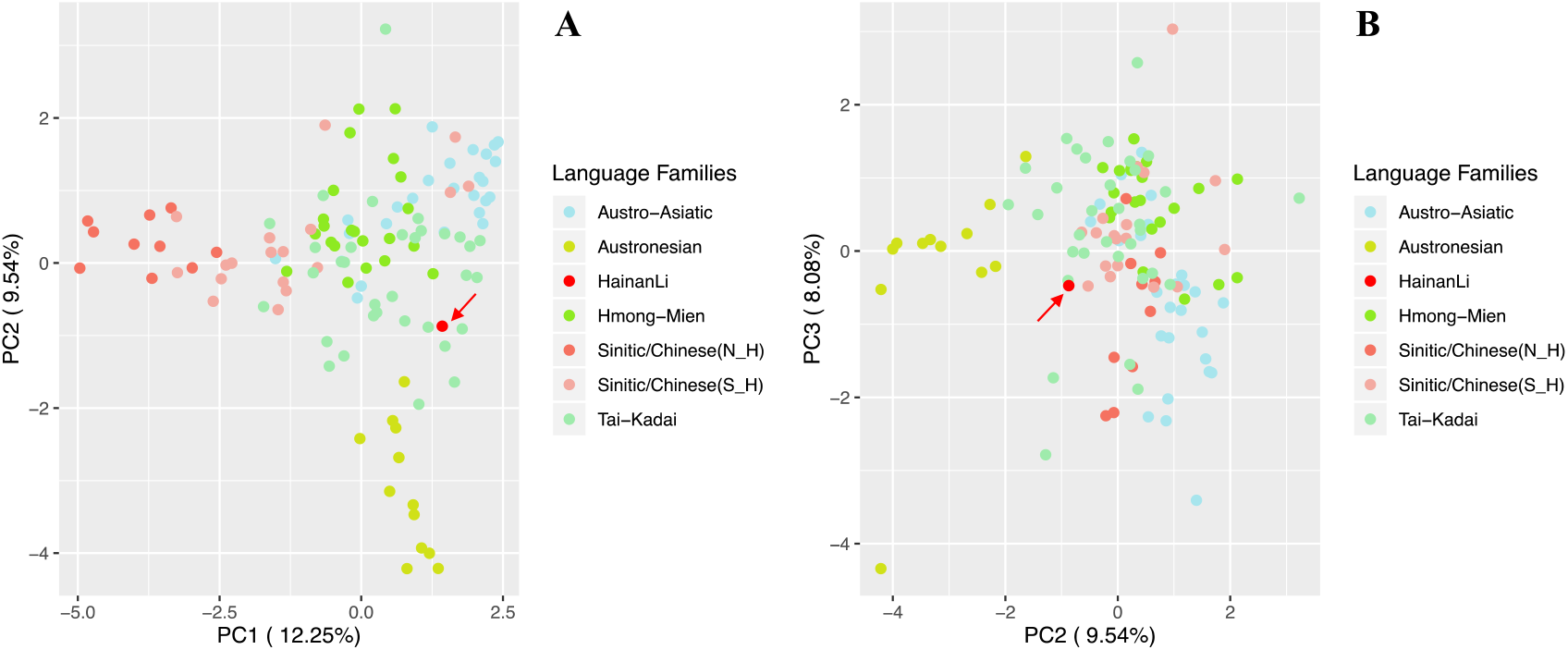
PCA based on Y-haplogroup frequencies in 123 populations of Dataset III (2). **A.** PCA (PC1 and PC2); **B.** PCA (PC2 and PC3).

The previous studies had illustrated that, the O1b1a1a-M95 lineage originated in the southern East Asia among the Tai-Kadai-speaking populations ~20-40 kya and then dispersed southward to Southeast Asia after the LGM before moving westward to the Indian subcontinent [183–186]. This haplogroup was shown to be prevalent in Austro-Asiatic speaking populations in Southeast Asia (74-87%) [179, 183, 187] and Northeast India (85%) [183, 187], the Tai-Kadai and Hmong-Mien speaking populations in China (45%) [12, 179–181, 188], and the Austronesian speaking populations (28%) [12, 183, 189, 190]. While, the O2-M122 lineage was dominant in East Asian populations, with an average frequency of 44.3% [164, 191]. The O2-M122 had the highest frequency in East Asians, especially in Han Chinese (52.06% in northern Han and 53.72% in southern Han), and was also quite frequent in Hmong-Mien (51.41%) and Austronesian (26.31%) populations, but it is absent outside East Asia. O2a2b1a1-M117 was another major founder paternal lineage [192, 193], comprising about 15-16% of modern Han Chinese [191, 192, 194] and was also quite frequent in Tibeto-Burman populations [188, 191, 195] like Nu (62%), Derung (32%), Lhoba (31%), Tibetan in Yunnan (22%) and Hani (17%). In short, the dominant lineage in the Hainan Li, O1b1a1a-M95 (45.40%), has a southern East Asia origin and are prevalent in indigenous people like Tai-Kadai populations, whereas the discovery of O2-M122 and O2a2b1a1-M117 in Hainan Li reflects that the limited gene flows existed between Hlai and Han Chinese (especially for the Southern Han) to some extent.

### 3.7 Discussions and significances

Above all, the MPS-based ForenSeq^™^ DNA Signature Prep Kit (DAPA, 152 forensically relevant genetic markers) was used for 136 Hainan Li individuals, and the related forensic landscape of Hainan Li was firstly presented in this study. The forensic characteristics and efficiencies of different types of genetic markers were evaluated by different bioinformatic methods and the phylogenetic analyses of the Hlai language-speaking population based on genetic profiles, linguistics and other aspects. The extended application of the ForenSeq^™^ DNA Signature Prep Kit (DAPA) based on the set of 94 iiSNPs can be conducted between the intercontinental populations for biogeography ancestry inference to some extent. It was proved that Hainan Li was an isolated population relatively on the basis of Dataset I and II, and had a close relationship with Hainan Lingao and a relatively far distance with Hainan Han. The analysis based on Dataset III further confirmed the limited gene flows in Hainan island, when the haplogroups O2-M122 and O2a2b1a1-M117 were observed in Hainan Li. From the perspective of linguistic, Hlai and Ong Be language belongs to Tai-Kadai language, while Hainanese is a branch of Sinitc/Chinese, no matter Tai-Kadai or Chinese, they all belong to Sino-Tibetan Language Family.

In past two millennia of Chinese history [9, 29, 30], there were three waves of large-scale migrations that took place during the Western Jin Dynasty (AD 265–316), the Tang Dynasty (AD 618–907), and the Southern Song Dynasty (AD 1127–1279), and many smaller southward migrations with continuous southward movements of Han people because of warfare, drought, and famine in the north, which accelerated that Han people and Han culture expanded into southern China, where inhabited by the southern natives initially, including those speaking Tai-Kadai, Austro-Asiatic and Hmong-Mien languages. The massive movement of the northern immigrants led to a change in genetic makeup in southern China, and resulted in the demographic expansion of Han people as well as their cultures [165]. Due to these population movements, gene flows between northern Hans, southern Hans and southern natives contributed to the admixture which shaped the genetic profiles of the extant populations.

Likewise, the current ethnic and demographic situations in southeast Asia influenced by three large scale immigrations since the Neolithic Age. The first population fusion occurred between the aboriginal Negritos and the Austronesian populations which had an origin-related controversial subject in linguistics and other related field from the West and East routes at BC 1,500-2,500 [196, 197]. The second movement to southeast Asia was the descent of the Mon-Khmer populations southward, leading to the movement to Indonesia and Philippines for the Austronesian 3,000 years ago [198, 199]. The migration of the Sino-Tibetan populations to the Indo-China Peninsula constituted the third large-scale event around 2,000 years ago [199–202]. The three massive population movement led to the in-depth exchanges of cultures and genetics between different migrations, and the migrations with the aboriginal Negritos, to take shape the mixture situation of multi-ethnics and multi-language in Southeast Asia nowadays [70].

Hainan Li, an isolated population, experienced limited gene flows with surrounding populations by virtue of geography, history and culture. Such populations include the Andaman Islanders [203, 204] and Sardinians [205–207] who maintained their unique allele frequency and phenotypic characteristics due to the geographic barriers; the Roma [178, 208, 209] and the Jews [210–212] have maintained genetic coherence over vast geographical distances because of their distinctive history and culture. Population isolation is more likely to generate population-specific haplotypes or lineages, allowing geneticists to trace population history. In addition, isolated populations are most helpful in some genetic studies. For example, the nosogenetic factors of the complex diseases are apparently reduced in the isolated populations and will be much easier to analyze [12]. Furthermore, with relatively big populations and long history, most of the Hainan aborigines have unlikely undergone genetic drift, and developed relatively high Y-STR diversity beside their low Y-SNP diversity. It is believed that Hainan aborigines could also have higher diversity of autosomal variances than the other East Asians, and lower linkage disequilibrium, which is more valuable for the disease association studies to avoid the false positive results caused by high linkage disequilibrium.

In forensic genetic fields, isolated populations can serve as stable models to construct the genetic structures for delicate population grids. The clarified population structures have benefits for biogeography ancestry inference for the partial and local subpopulations. Furthermore, the unique and slight Y haplogroup, because of gene flows with surrounding populations, delivers another potential clue for forensic application to some extent. Back to this study, the DNA profiles of the MPS-based ForenSeq^™^ DNA Signature Prep Kit (DAPA) in Hainan Li were reported for the first time, which characterized the forensic polymorphism landscape. The sequence-based genetic markers featured more genetic diversity of loci and improved the system efficiency, more forensic associated studies based on MPS technology and related products could be conducted widely, which could strengthen the high-throughput application in forensic science and establish compact connections between the length-based and the sequence-based data as soon as possible.

## 4. Conclusion

In summary, the forensic landscape of genetic polymorphism data of 136 Hainan Li by genotyping 152 forensic-associated genetic markers using the ForenSeq^™^ DNA Signature Prep Kit (DAPA) were characterized for the first time in this study. Allele frequencies based on the diversities of both length and sequence were reported, and the forensic parameters were evaluated comprehensively from different genetic scales. The results demonstrated that the STRs and SNPs analyzed here were highly polymorphic in Hainan Li population and could be employed in forensic applications, in addition, the set of SNPs, because of the allele frequencies disequilibrium in some loci, expanded potential application for biogeography ancestry inference between the intercontinental populations. The phylogenetic analyses indicated that Hainan Li, with a southern East Asia origin and Tai-Kadai language-speaking language, is an isolated population relatively, and has a close relationship with Hainan Lingao, and a relatively far distance with Hainan Han in isolated Hainan island, but the role of the limited gene flows from the Tai-Kadai populations and Hainan Han in shaping the genetic pool of Hainan Li can’t be ignored. In addition, the isolated population models will be beneficial to clarify the exquisite population structures and develop specific genetic markers for subpopulations in forensic genetic applications.

## Acknowledgements

We would like to thank all the volunteers who contributed samples for this study. Especially, I’d like to express the sincere gratitude to *Miss Lijuan Yang* for the image processing of Fig. 1. This study was supported by the Program of Hainan Association for Science and Technology Plans to Youth R&D Innovation (QCXM201705), the National Undergraduate on Innovation and Entrepreneurship Training Program (201911810008 and 201911810023), and the National Natural Science Foundation of China (No.81560304).

