## Supplemental Figures 1-6 for "The forensic landscape and the population genetic analyses of Hainan Li based on massively parallel sequencing DNA profiling"

### Slide 1
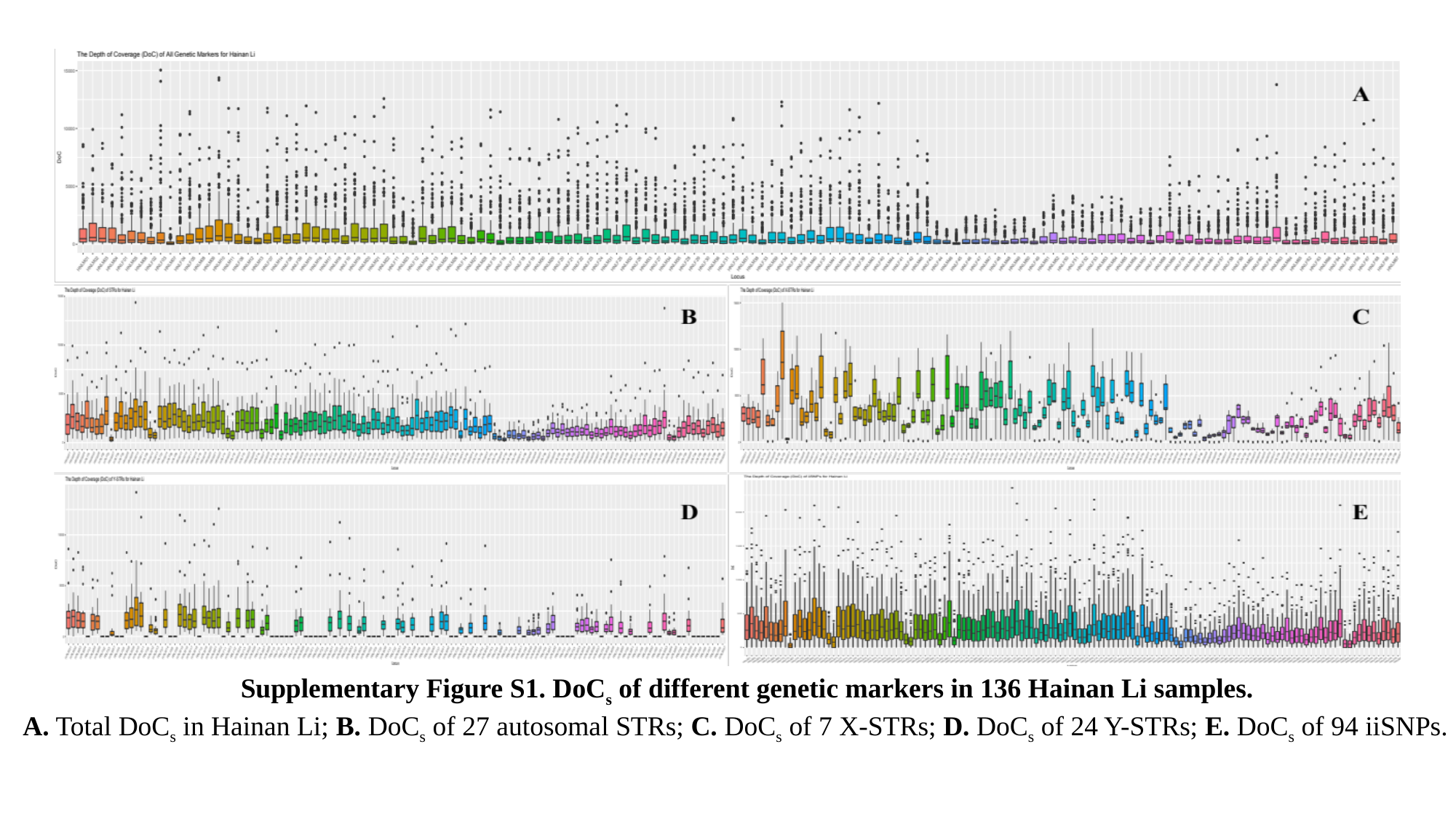

Supplementary Figure S1. DoC_s_ of different genetic markers in 136 Hainan Li samples.
A. Total DoC_s_ in Hainan Li; B. DoC_s_ of 27 autosomal STRs; C. DoC_s_ of 7 X-STRs; D. DoC_s_ of 24 Y-STRs; E. DoC_s_ of 94 iiSNPs.

### Slide 2
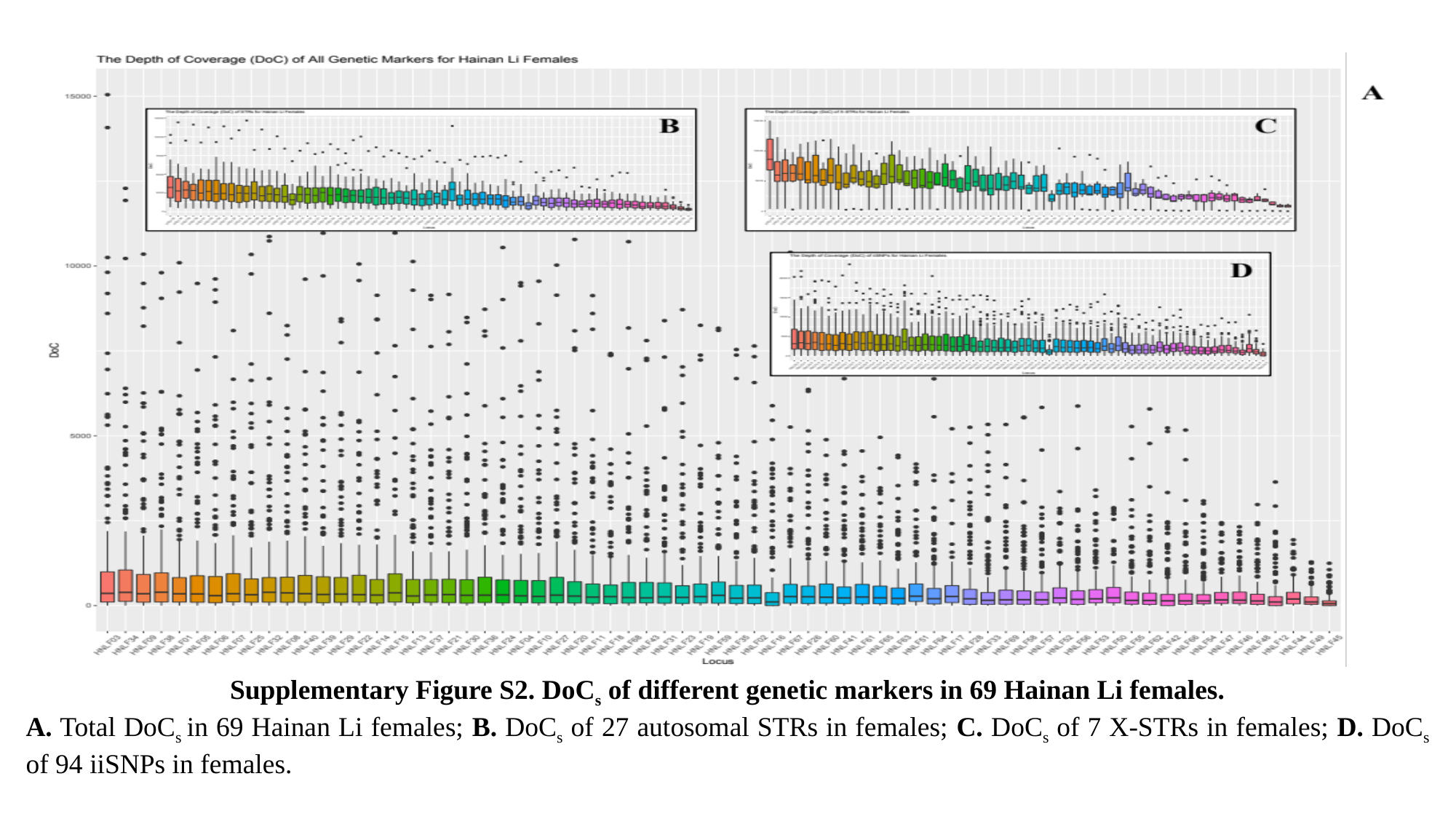

Supplementary Figure S2. DoC_s_ of different genetic markers in 69 Hainan Li females.
A. Total DoC_s_ in 69 Hainan Li females; B. DoC_s_ of 27 autosomal STRs in females; C. DoC_s_ of 7 X-STRs in females; D. DoC_s_ of 94 iiSNPs in females.

### Slide 3
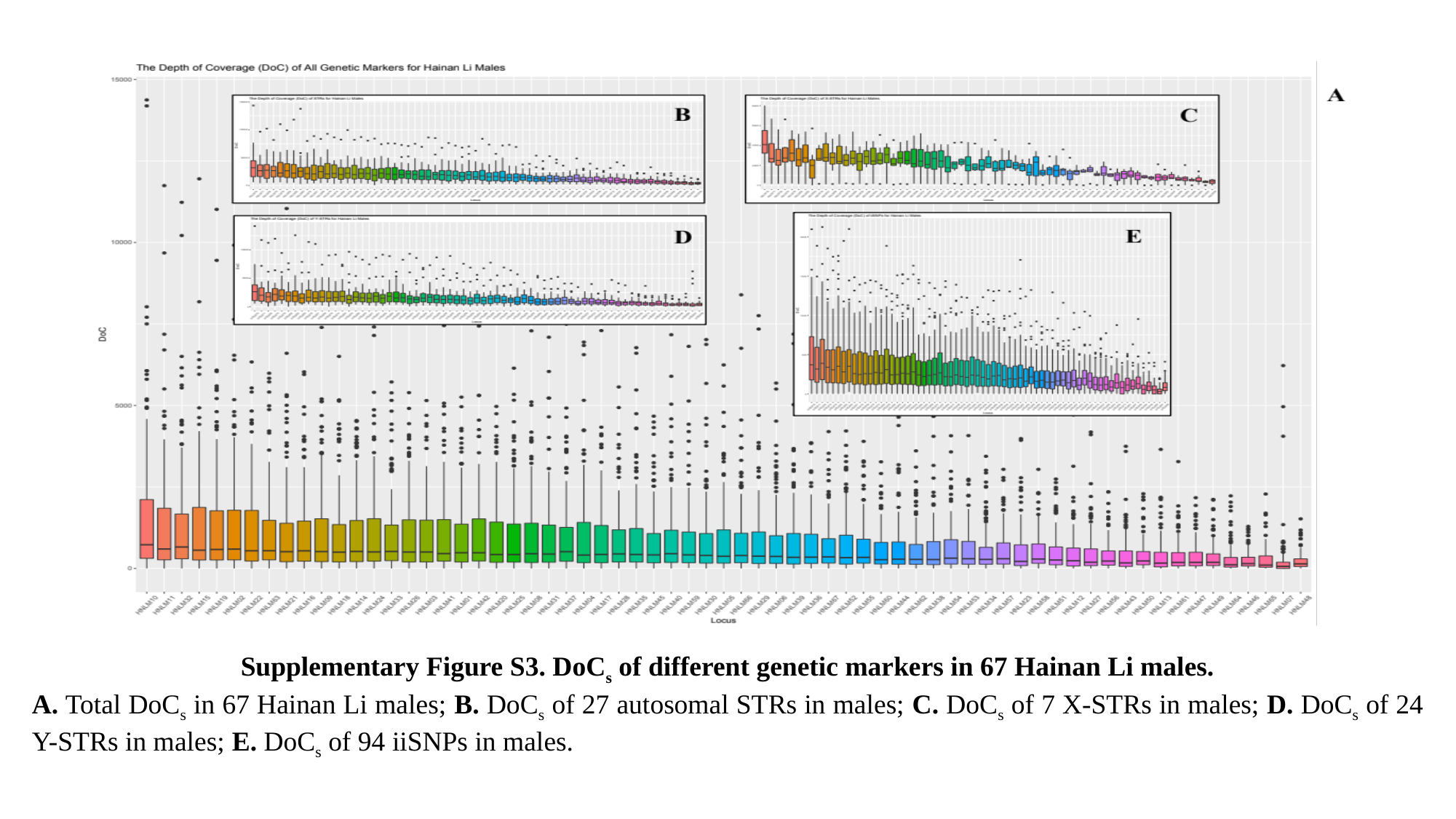

Supplementary Figure S3. DoC_s_ of different genetic markers in 67 Hainan Li males.
A. Total DoC_s_ in 67 Hainan Li males; B. DoC_s_ of 27 autosomal STRs in males; C. DoC_s_ of 7 X-STRs in males; D. DoC_s_ of 24 Y-STRs in males; E. DoC_s_ of 94 iiSNPs in males.

### Slide 4
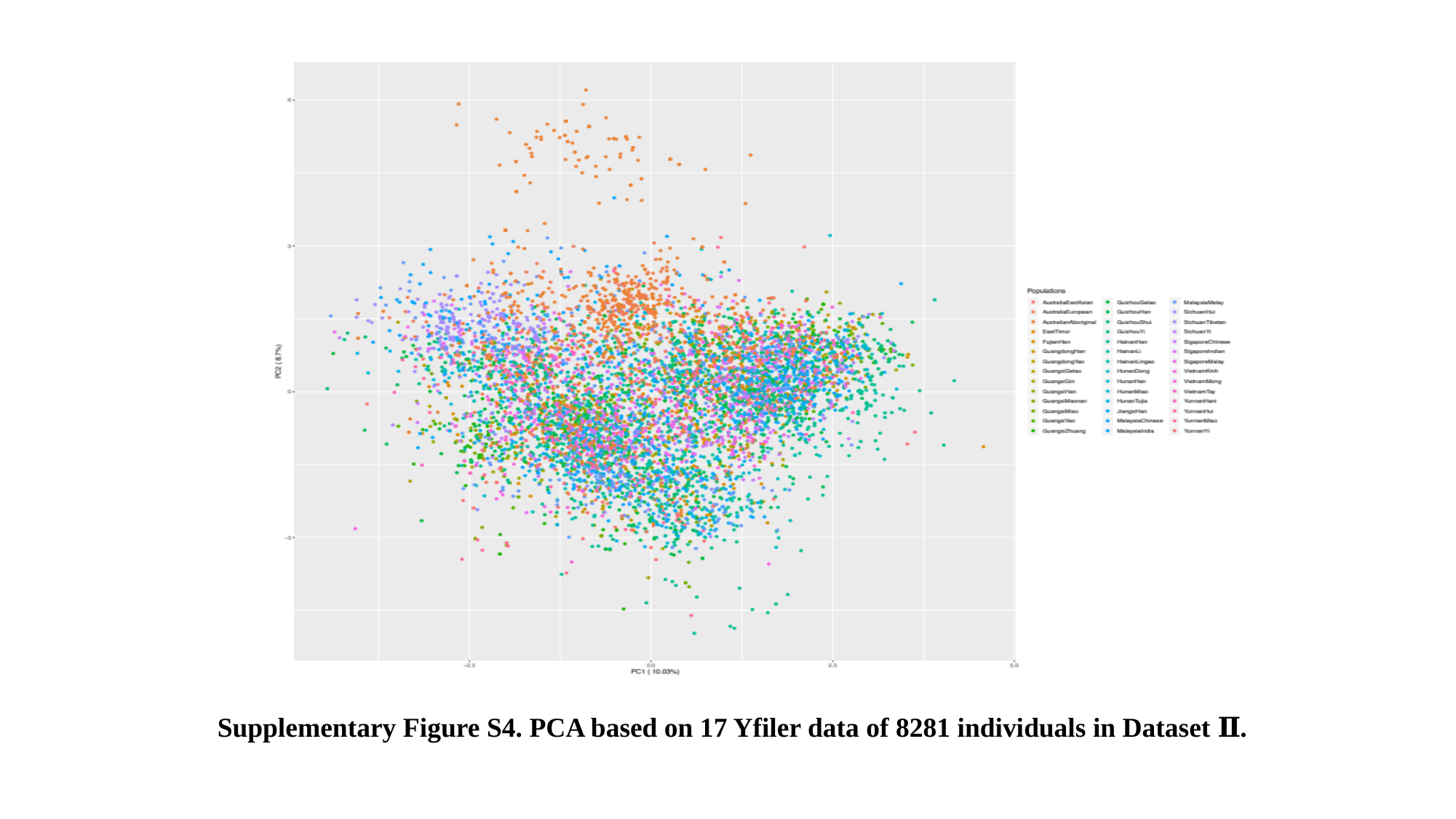

Supplementary Figure S4. PCA based on 17 Yfiler data of 8281 individuals in Dataset Ⅱ.

### Slide 5
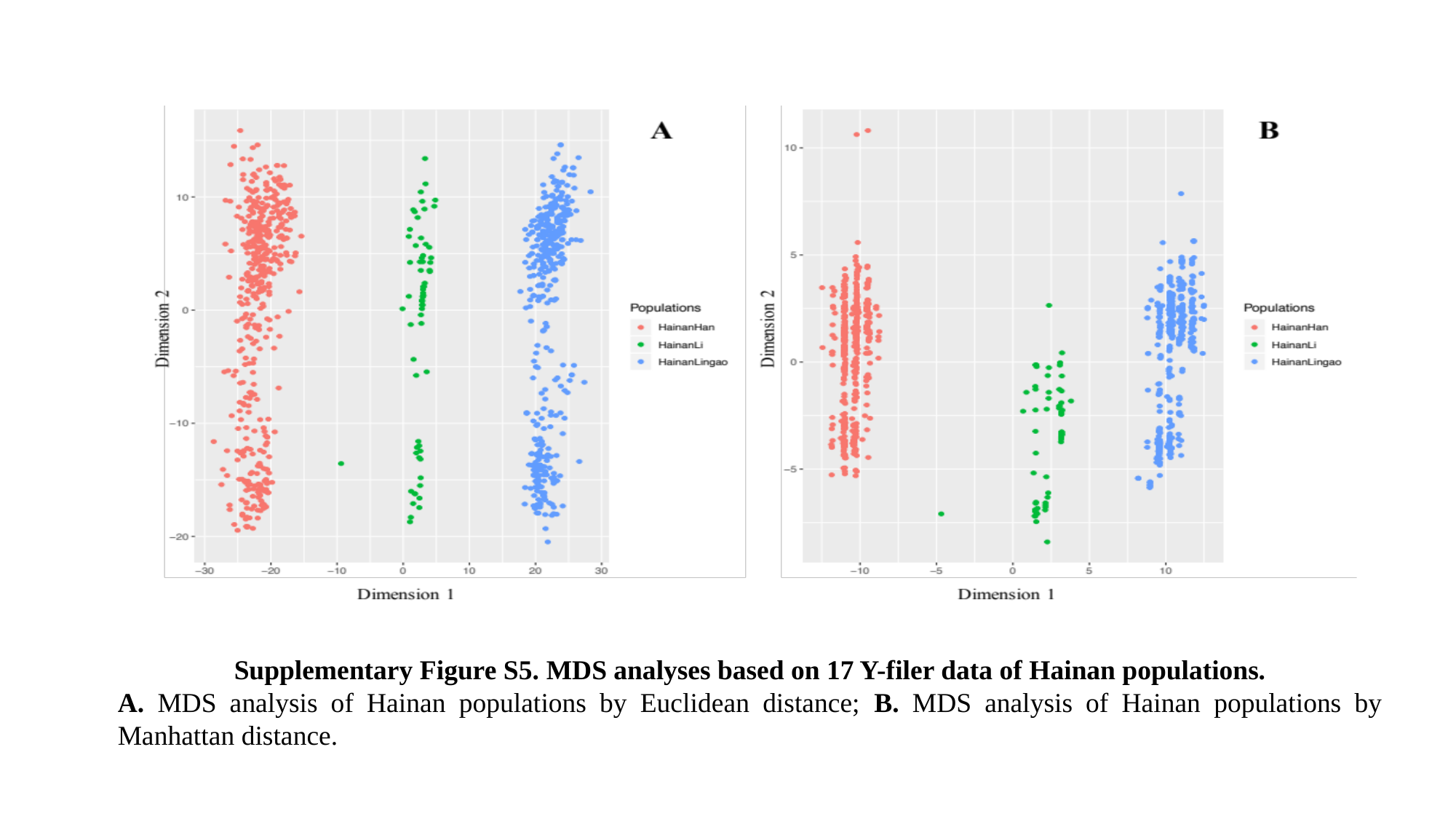

Supplementary Figure S5. MDS analyses based on 17 Y-filer data of Hainan populations.
A. MDS analysis of Hainan populations by Euclidean distance; B. MDS analysis of Hainan populations by Manhattan distance.

### Slide 6
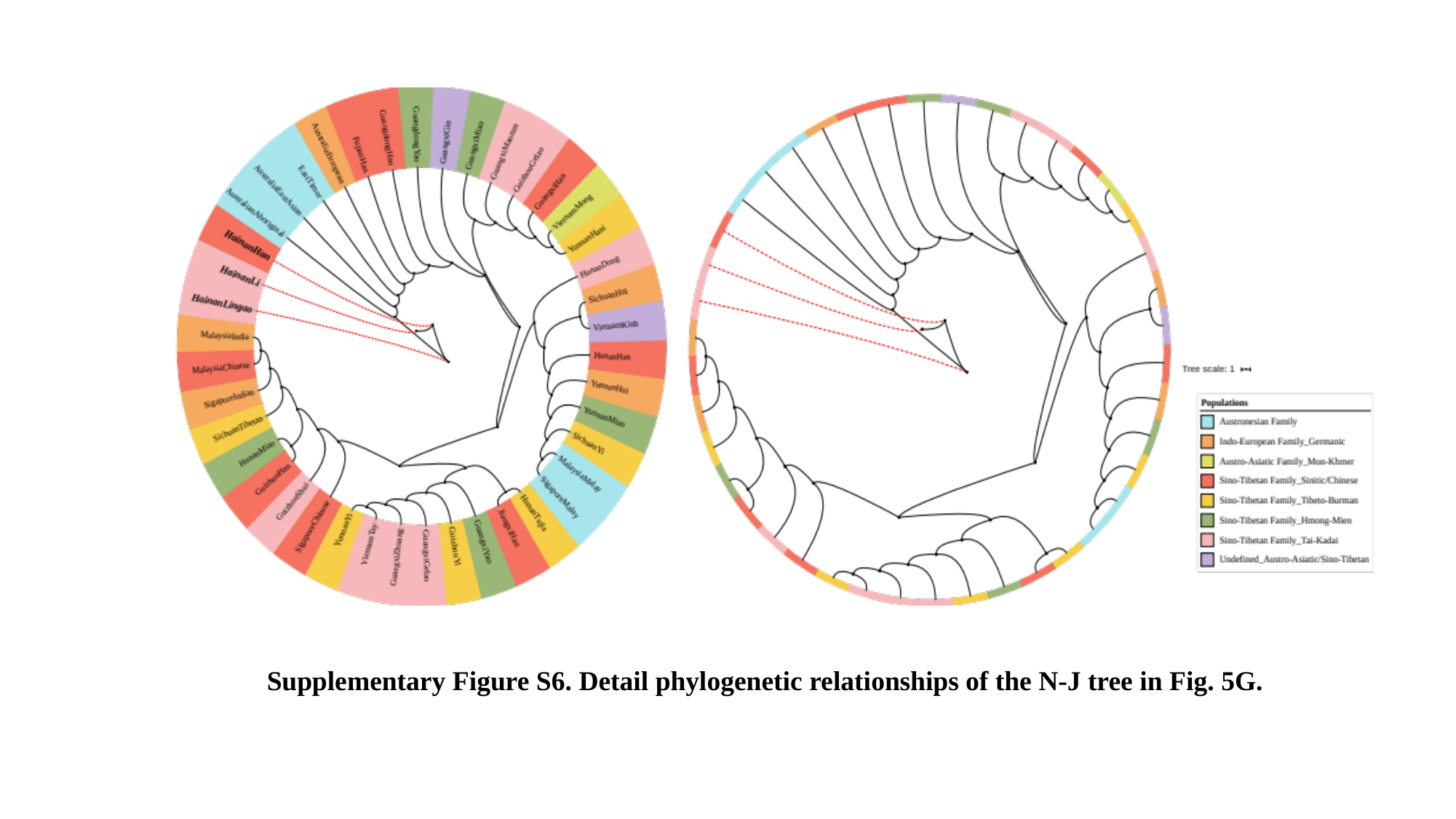

Supplementary Figure S6. Detail phylogenetic relationships of the N-J tree in Fig. 5G.
